# Hyper-responsiveness of cancer stem cells to microenvironmental cues controls metastasis and therapy response through YAP/TAZ/TEAD

**DOI:** 10.1101/2025.03.13.643008

**Authors:** Binwu Tang, Jacob Minin, Victoria Gonzalez, Zoya Khan, Yuval Raviv, Yu-an Yang, Christine Carney, Zachary Millman, Daniel Grun, Cristiana Pineda, Alina Sharma, Dominic Esposito, Hualong Yan, Jing Huang, Andy D. Tran, Michael Kruhlak, Howard Yang, Maxwell P. Lee, Lalage M. Wakefield

## Abstract

Cancer stem cells (CSCs) are key drivers of metastasis and therapy resistance but have been challenging to visualize and study *in situ*. Using a fluorescent CSC reporter, we observed very different population dynamics for CSCs and nonCSCs during metastatic lung colonization in breast cancer models. CSC expansive self-renewal drives early lesion formation before switching to a maintenance mode of balanced self-renewal and differentiation, whereupon nonCSC proliferation takes over as the main driver of metastatic expansion. Mechanistic analyses showed that CSCs are hyper-responsive to microenvironmental cues such as cell crowding and nutrient availability, suggesting a novel role for CSCs as sensors and early responders to fluctuating local conditions in the tumor. Incoming signals converge on YAP/TAZ/TEAD, with heightened CSC sensitivity and response supported by elevated receptor expression and increased chromatin accessibility around enhancers with TEAD binding sites. Targeting inputs to the YAP/TAZ/TEAD node reversed chemotherapy-induced enrichment of CSCs in lung metastases.

**Highlights:**

- Different population dynamics for breast cancer stem cells (CSCs) and their differentiated progeny in early metastatic colonization
- CSCs are hyper-responsive to microenvironmental cues and serve as sensors of local conditions for the tumor
- Many microenvironmental inputs converge on YAP/TAZ to regulate self-renewal vs differentiation fate decisions in the CSC
- Targeting YAP/TAZ input pathways blocks chemotherapy-induced enrichment of CSCs

## INTRODUCTION

Many epithelial cancers are initiated and sustained by a small subpopulation of tumor cells with stem-like properties, the “cancer stem cells” (CSCs)^1, 2^. CSCs are uniquely capable of indefinite self-renewal and give rise to the more differentiated progeny that form the bulk of the tumor. Metastasis is driven by “metastasis-initiating cells”, which likely represent a subset of CSCs with acquired features that allow them to intravasate and survive the stresses encountered during dissemination and distant colonization^3,4, 5^. CSCs are often phenotypically similar to fetal somatic stem cells or facultative stem cells that are recruited for tissue repair^3, 6, 7^. Thus fetal or regenerative stem programs may be co-opted by oncogenically-initiated cells to generate CSCs. The clinical significance of CSCs lies in their relative resistance to nearly all forms of therapy^1, 8, 9^. This resistance results from intrinsic properties such as elevated drug efflux pumps and DNA repair mechanisms, and from the generation of protective niches^8, 10, 11^. Consequently, CSCs drive therapeutic relapse and disease recurrence. A better understanding of CSC dynamics will be critical for development of more effective therapeutics.

Like somatic stem cells, CSCs exist on a phenotypic continuum that has the most “stem-like” cells (CSCs) at one end and the most differentiated cells at the other^12^. However, phenotypic plasticity is higher in tumors than in normal tissues, enabling some bidirectional movement along this continuum rather than the normal unidirectional differentiation^1, 12^. The study of CSCs has been inherently challenging due to this plasticity, their small numbers, and the shortage of consensus markers for their identification *in situ*. To address these challenges, we developed a lentiviral-based fluorescent CSC reporter that reports on stemness as a dynamic phenotypic state, with good temporal resolution for live-cell imaging *in vitro* and *in vivo*^13, 14^. Intravital imaging with this reporter in breast cancer models previously revealed new dynamic attributes of CSCs in primary tumors *in vivo*, including the ability of macrophage contact to confer stem properties on more differentiated tumor cells (nonCSCs) ^14^.

In the current study, we used this CSC reporter in breast cancer models to generate detailed information on the population dynamics of CSCs and nonCSCs during early metastatic colonization of the lung. Modeling these dynamics *in vitro*, we uncovered a novel role for CSCs as hyper-responsive microenvironmental sensors for the tumor cell population as a whole. Signals from microenvironmental cues are integrated at the YAP/TAZ/TEAD transcriptional hub, and we show that the CSC hyper-responsiveness is due to increased cell surface receptors for these input signals, as well as more open chromatin at gene loci with TEAD binding sites in their enhancers. Importantly, blockade of upstream inputs to the YAP/TAZ/TEAD hub reversed the chemotherapy-induced enrichment of CSCs in treated metastases.

## RESULTS

### Breast cancer stem cells are enriched in the circulation and on arrival in the lung

Our fluorescent lentiviral CSC reporter (SORE6>FP) dynamically reports on stemness through activation of an artificial enhancer (“SORE6”) by the stemness master transcription factors, Oct4 and Sox2 or their paralogs^13, 14^. A control construct lacking the SORE6 enhancer but retaining the minimal CMV promoter (minCMV) is used for gating or thresholding (**Fig. 1a, Extended Data Fig. 1a**). The destabilized mCherry used here has a half-life of ∼1.4h (**Extended Data Fig. 1b**), giving the reporter high selectivity and temporal resolution. For this study, breast cancer cell lines were transduced with a constitutive nuclear GFP marker (all tumor cell nuclei green), and either SORE6>dsmCherry (CSCs marked red/yellow) or the matched control construct. Representative flow profiles for transduced tumor cells are given in **Extended Data Fig. 1c**. FACS-sorted SORE6+ cells were significantly enriched for tumor-initiating ability by *in vivo* limiting dilution analysis in all models tested (**Extended Data Fig. 1d,e**), so SORE6+ cells will be referred to as CSCs in our subsequent analyses. In breast cancer models, our reporter enriches both tumor-initiating and metastasis-initiating activity in the Oct4^hi^ Sox2^hi^ SORE6+ population, and we use the umbrella term CSC throughout.

**Fig. 1.**
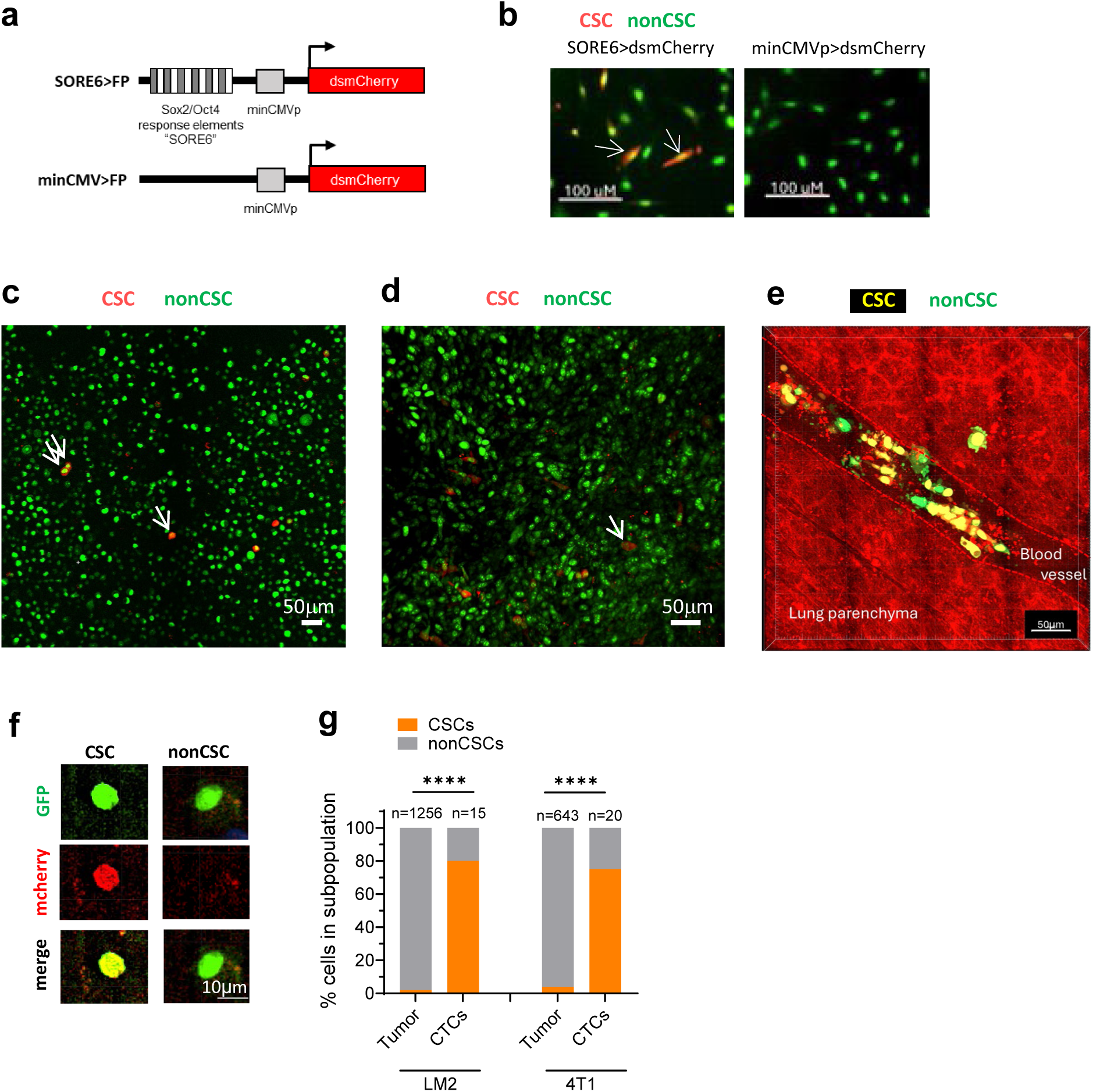
| Breast cancer stem cells are enriched in the circulation and on arrival in the lung. **a.** Schematic of SORE6 cancer stem cell and matched control reporter. minCMVp is a minimal CMV promoter. **b.** Epifluorescent images of MDAMB231-LM2 cells transduced with the SORE6>dsmCherry reporter or minCMV>dsmCherry control construct growing *in vitro*. All tumor cells have a constitutive nuclear GFP marker. Arrows indicate CSCs. **c,d** Representative confocal images of fresh orthotopic primary tumors from LM2 model **(c)** and 4T1 model **(d)**. Arrows indicate CSCs; double arrow indicates rare CSC doublet **e.** High resolution image of LM2 cells arriving in the lung 10 weeks after orthotopic implantation of a primary tumor. **f.** Representative image of circulating tumor cells from a mouse with an LM2 primary tumor **g.** Quantitation of % CSCs in primary tumor and circulating tumor cells (CTCs) from LM2 or 4T1 primary tumors. Fisher Exact test. Number of cells (n) assessed is indicated for each condition.

To address CSC dynamics during the early stages of triple negative breast cancer (TNBC) metastasis to the lung, we used the human MDAMB231-LM2 (hereafter “LM2”)^15^ and the mouse 4T1 cell line models^16^. The SORE6 reporter marked a minority population of LM2 cells *in vitro* (**Fig. 1b**), and in orthotopic primary tumors *in vivo* (**Fig. 1c**). Similar results were seen for 4T1 primary tumors (**Fig. 1d**). CSCs typically appeared as isolated single cells in large primary tumors, with only occasional doublets, usually towards the tumor edge. In **Fig. 1e**, we captured several tumor cells from an LM2 orthotopic tumor simultaneously arriving in the vasculature at the lung metastatic site. 25 of the 32 tumor cells (78%) in the field were CSCs, suggesting a strong enrichment of CSCs among intravasated tumor cells. Indeed, we found that circulating tumor cells recovered from the blood of mice with LM2 or 4T1 orthotopic tumors were 75-80% CSCs, compared with 1-2% CSCs in the primary tumors (**Fig. 1f,g**) confirming the enrichment of CSCs in the circulation that has been observed previously in clinical and preclinical settings^17,14^. Thus this powerful imaging tool is performing as expected and enables quantitative insights into CSC and nonCSC dynamics during tumor progression.

### Different population dynamics for CSCs and nonCSCs in early lung colonization

We next undertook a detailed analysis of CSC dynamics following tumor cell arrival in the lung. To generate enough metastatic events for quantitative analysis, we increased metastatic efficiency for the LM2 model by using a pseudo-orthotopic format with tail vein injection, while the 4T1 model involved orthotopic implantation and tumor resection (**Fig. 2a**). Unless otherwise noted, unsorted tumor cell cultures containing both CSCs and nonCSCs were injected. CSCs and nonCSCs in metastatic lesions were quantitated by *ex vivo* imaging of freshly excised lungs at different times. For the LM2 model, nonCSCs died off rapidly within two days after arrival in the lung, while CSCs mostly maintained their numbers, such that by Day2 the % CSCs in the lung was similar to that seen arriving in the lung from orthotopically implanted tumors (**Extended Data Fig. 2a**, compare with **Fig. 1e,g)**. In all subsequent tail vein injection experiments, we started analysis at Day 3 after this initial tumor cell die-off, to better mimic the situation seen in the more realistic orthotopic format.

**Fig. 2.**
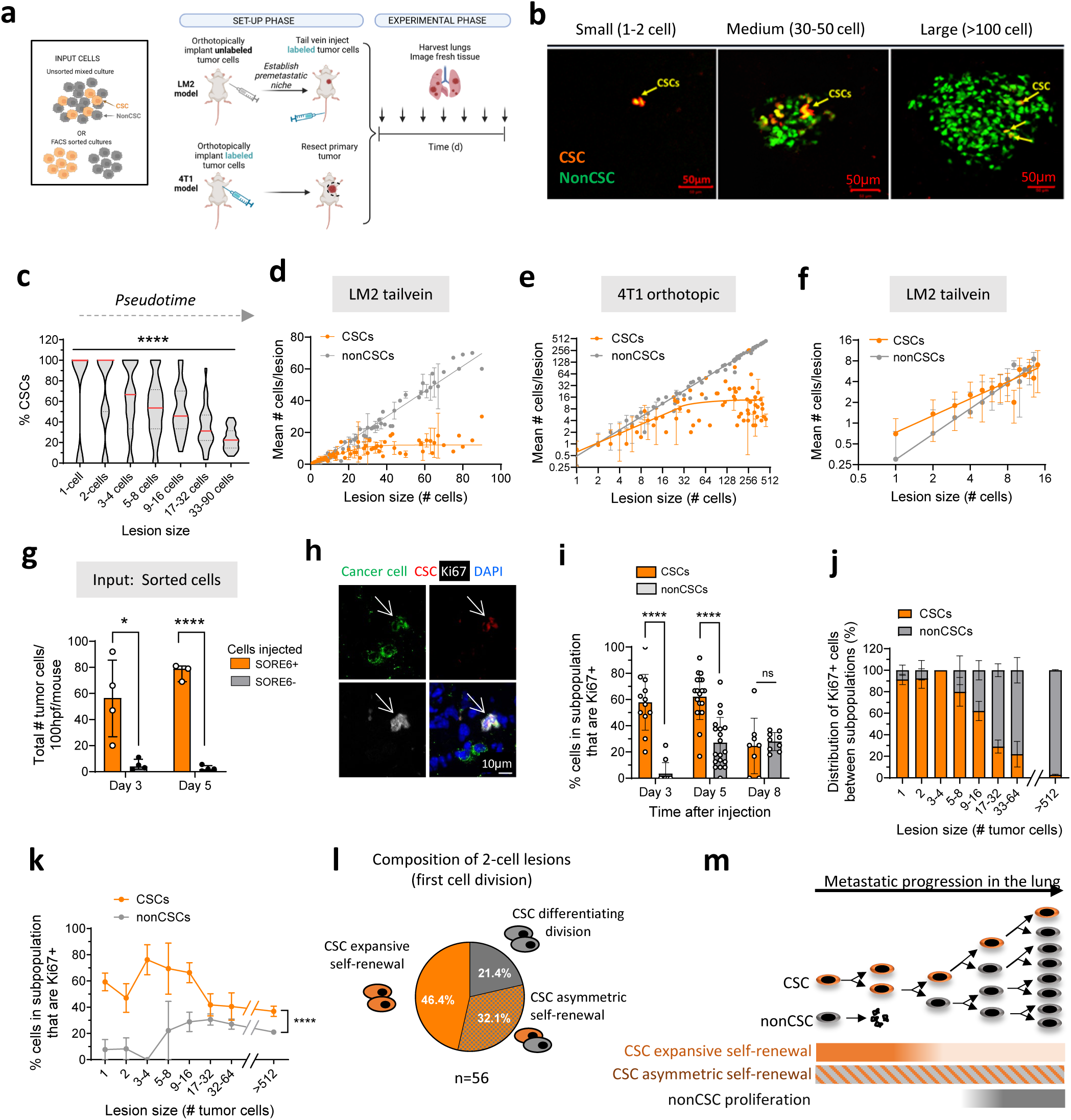
| CSCs and nonCSCs have different population dynamics in early lung colonization. **a.** Schematic for lung metastasis study formats for LM2 and 4T1 models. **b.** Representative images of LM2 lung metastatic lesions of different sizes. **c.** CSCs as % total tumor cells in LM2 metastatic lesions of different sizes. Data pooled from multiple timepoints for a total of 26-88 lesions analyzed/size bin from 14 mice. Red bars show medians. One-way ANOVA. **d**. Different kinetics for expansion of LM2 CSC and nonCSC populations in the lung as a function of lesion size following tail-vein injection. Results are mean +/− SD for n=1-63 depending on lesion size (pooled from 14 mice and 512 lesions over multiple timepoints). **e.** Expansion kinetics for CSC and nonCSC populations in the lung of mice with orthotopic 4T1 tumors. Results are mean +/− SD where n=1-29 depending on lesion size (pooled from n=9 mice and n=148 lesions over multiple timepoints). Note log2 axes. **f.** Logplot showing detailed expansion kinetics of LM2 metastases at small lesion sizes. Results are mean +/− SD for n= 1-29 lesions/size bin. **g.** Tumor cells in lung from mice injected with sorted SORE6+ or SORE6-LM2 cells by tail-vein injection. Results are median +/− IQ range, n=3-4 mice/group, unpaired t-test **h.** LM2 lung metastases immunostained for Ki67 (proliferation), GFP (tumor cells), mCherry (CSCs). Arrow indicates CSC. **i.** % Ki67+ proliferating cells within CSC or nonCSC subpopulations after injection of unsorted cells. Results are median +/− IQ range, 8-24 total lesions/timepoint, Mann-Whitney test. **j.** Distribution of proliferating cells between CSC and nonCSC compartments as a function of LM2 metastatic lesion size (mean +/− SEM; n=3-35 lesions/size bin; data for largest size bin is from an independent expt (n=15 lesions)). **k.** % cells within CSC or nonCSC subpopulations that are Ki67+ in LM2 lung lesions of different sizes. Results are mean +/− SEM, n=3-54/size bin; data for largest size bin from an independent expt (n=15 metastases; Mann-Whitney U test) **l.** Composition of 2-cell lesions in LM2 model (n=56 lesions from 2 expts). **m.** Model for CSC and nonCSC kinetics in early metastatic colonization.

As expected, LM2 cells formed progressively growing metastatic lesions (**Extended Data Fig. 2b**). The overall % CSCs among tumor cells in the lung decreased with time after implantation (**Extended Data Fig. 2c**). However by later times, a large range of lesion sizes co-exist in the same lung (**Extended Data Fig. 2b**), with day 14 lesions ranging from 1 cell to >300 cells. A major advantage of our imaging approach is that the number of CSCs can be assessed for individual metastatic lesions, allowing for a more accurate assessment of cellular dynamics during lesion expansion. The CSC/nonCSC composition of lesions of a given size was very similar regardless of the time of lung harvest (**Extended Data Fig. 2d**). This suggests that there may be a variable latency period for outgrowth of single tumor cells after lung seeding, but that once outgrowth begins, individual lesions show similar population dynamics. Thus, we can pool lesions from different timepoints within an experiment, using lesion size as a measure of pseudotime. With this approach, we found that the % CSCs decreased as metastatic lesion size increased (**Fig. 2b,c**). Quantitating absolute numbers of CSCs and nonCSCs in individual lesions as a function of lesion size revealed complex population dynamics. The CSC subpopulation reached an early plateau when the metastatic lesion size was ∼32 cells total, while the nonCSC subpopulation expanded continuously (**Fig. 2d**). Similar kinetics were observed for LM2 metastasis from the orthotopic site, though the number of evaluable lesions was much smaller (**Extended Data Fig. 2e).** The orthotopic mouse 4T1 model also showed similar kinetics (**Fig. 2e, Extended Data Fig. 2f**).

Looking in detail at the smallest lesions in the LM2 model, we were surprised to find that the nonCSC subpopulation expands faster than the CSC subpopulation (**Fig. 2f**). This finding was consistent across models and formats (**Extended Data Fig. 2g,h**), raising the unexpected possibility that early metastatic colonization is driven by proliferation of the nonCSC subpopulation. This would challenge the notion that only CSCs can initiate metastases. However, injecting sorted SORE6+ CSCs or SORE6-nonCSCs from the LM2 or 4T1 models confirmed that nonCSCs cannot generate progressively growing lesions (**Fig. 2g; Extended Data Fig. 2i**). To resolve this apparent discrepancy, we assessed proliferation in both subpopulations by immunostaining for Ki67 (**Fig. 2h**). At early times (day 3-5 after injection), the majority (∼60%) of the CSCs in metastatic lesions were actively proliferating (**Fig. 2i**). In contrast, very few nonCSCs were in cycle at day 3, and only began to proliferate around day 5. Plotting the data as a function of lesion size confirmed that the majority of proliferating cells in smaller lesions are CSCs, while nonCSCs are the majority of proliferating cells in larger lesions (**Fig. 2j**). Importantly however, even in the largest lesions, the CSCs as a class are still significantly more proliferative than the nonCSCs **(Fig. 2k)**.

Collectively, these data suggest that early expansion of the nonCSC subpopulation in metastases must be driven by differentiation of CSCs and not by proliferation of nonCSCs. Analysis of 2-cell lesions, representing the first cell division, shows that 29% have one CSC and one nonCSC, consistent with nearly a third of the early CSC divisions being asymmetric (**Fig. 2l**). Another 23% of 2-cell lesions had no CSCs, suggesting that some CSCs may also undergo fully differentiating divisions. However, it is worth noting that we rarely see lesions of >32 cells without a CSC (**Extended Data Fig. 2j**), suggesting that if 2-cell lesions without CSCs progress, they either exhaust their proliferative capacity within ∼5 doublings, or must regenerate CSCs to expand further.

Overall, our data lead to a model for early metastasis dynamics (**Fig. 2m**). Following arrival of tumor cells in the lung, most nonCSCs and some CSCs die off. Initiation of progressively growing metastatic lesions is driven solely by proliferation of CSCs. This CSC proliferation is a mix of symmetric self-renewing divisions, leading to expansion of the CSC population, as well as asymmetric divisions which give rise to a CSC and a nonCSC. At about the 32-cell stage, the CSC population ceases to expand, although ∼40% of the CSCs are still proliferating. Thus CSC divisions in larger lesions must be predominantly asymmetric, either physically or statistically. These asymmetric divisions maintain CSC numbers and refresh the nonCSC population. Expansion of larger lesions is driven primarily by proliferation of the much more numerous nonCSC progeny.

### Early inhibition of CSC population expansion is density-dependent and driven by increased CSC differentiation

The *in vivo* data showing an early CSC plateau suggests there is a negative feedback mechanism in larger metastases that acts on CSC fate decisions. Understanding this mechanism might help identify targetable pathways to selectively eliminate CSCs. Using Incucyte® Live Cell imaging for simultaneous real-time monitoring of CSCs and nonCSCs (**Fig. 3a**), we found that the early shut-down of CSC expansion in LM2 cells could be modeled *in vitro* (**Fig. 3b**). With time in culture, the CSC population reached an early plateau, while the nonCSC population continued to expand. Consequently the % CSCs in the culture is continually changing (dotted blue line). Broadly similar kinetics were seen for other TNBC models and the Ras-transformed mammary epithelial line MCF10Ca1h (**Fig. 3c-g**), though in some cases the CSC peak was followed by a loss phase rather than a plateau.

**Fig. 3.**
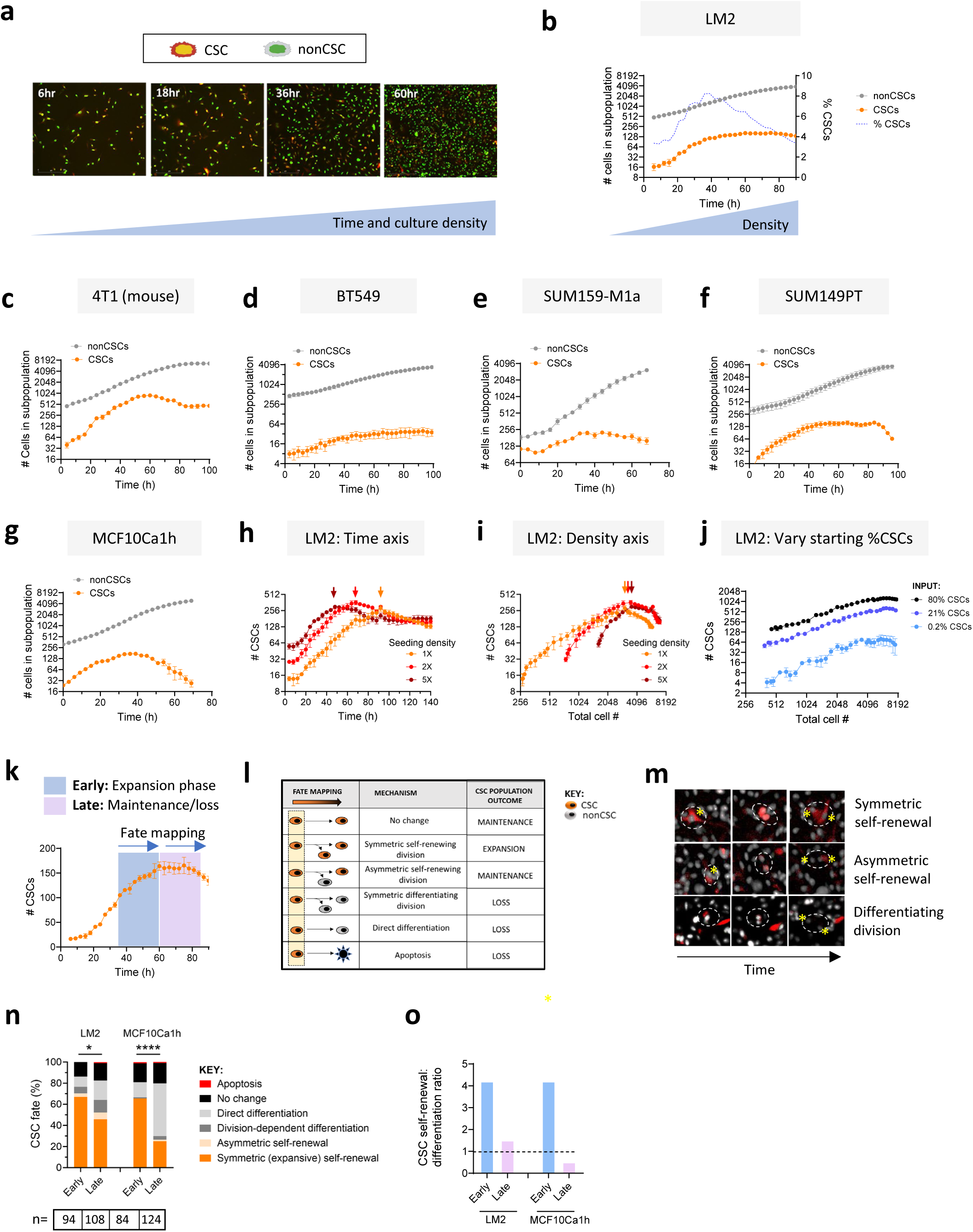
| Early inhibition of CSC population expansion is density-dependent and driven by increased CSC differentiation. **a.** CSCs and nonCSCs in unsorted LM2 cultures *in vitro* with increasing time in culture. **b.** Expansion kinetics for CSCs and nonCSCs in unsorted LM2 cells *in vitro* as a function of time from seeding. The % CSCs is shown on the right y-axis. Results are mean +/− SD, n=3. **c-g** Expansion kinetics of CSCs and nonCSCs in unsorted cultures of additional breast cancer cell lines as indicated. **h.** Effect of initial seeding density on LM2 CSC expansion kinetics as a function of time in unsorted cultures. NonCSCs are not shown. Mean +/− SD, n=3/timepoint. **i.** Data from (**h**) replotted as a function of culture density (total CSCs + nonCSCs). **j.** CSC expansion kinetics as a function of culture density for FACS-sorted LM2 cell populations with different starting % CSCs. Mean +/− SD, n=3 **k.** Time-frame for fate mapping of CSCs in relation to their expansion and maintenance/loss phase in unsorted LM2 cultures. **l.** Schematic for all possible CSC cell fates in fate-mapping experiments. **m.** Representative time-lapse images showing different types of CSC cell divisions. Red: cytoplasmic mCherry (CSCs); White: nuclear GFP (all tumor cells). Yellow asterisks indicate parent CSC and mitotic daughters. **n.** CSC cell fates during expansion (early) phase and maintenance/loss phase (late) for LM2 and MCF10Ca1h cells. Fisher Exact test for self-renewal vs other fates. **o.** Ratio of self-renewing divisions to differentiating events (direct or division-dependent) in LM2 and MCF10CA1h cells. Data from (**n)**.

With higher initial seeding densities, the CSC population peaked earlier (**Fig. 3h**). However, when CSCs were plotted as a function of the number of total cells (CSCs plus nonCSCs) in the culture, the CSC population size always peaked at the same cell density (**Fig. 3i**). Our initial hypothesis was that once the nonCSC population reaches a critical size, it might feedback inhibit the CSCs. However, when we varied the % CSCs in the input culture from 0.2% to 80%, we found that CSC numbers plateaued at the same total cell density regardless of the composition of the culture at the plateau stage (**Fig. 3j, Extended Data Fig. 3a**). Thus we reasoned that the CSC plateau is driven by some change in the culture that is independent of its cellular makeup at the critical culture density.

To address what happens to CSCs at higher cell density, we performed single cell fate mapping of CSCs over the time intervals indicated in **Fig. 3k**. Possible CSC fates are schematized in **Fig. 3l**, with representative images in **Fig. 3m**. We found a significant increase in the frequency of CSC differentiation, with a concomitant reduction in expansive self-renewal, in the later CSC plateau/loss phase compared with the earlier expansion phase (**Fig. 3n**). The reduction in the CSC self-renewal to differentiation ratio was more pronounced in cell lines such as MCF10Ca1h which showed loss of CSCs at later times (**Fig. 3n,o**). Consistent with our *in vivo* data, most CSCs were still proliferating at the CSC plateau in the LM2 model (**Extended Data Fig. 3b)**. In contrast, MCF10Ca1h CSC numbers declined after their peak due to a decrease in proliferation, and direct (division-independent) differentiation was the major CSC fate at later times. Thus increasing culture density leads to a change in cell fate decision in the CSC subpopulation, tipping the balance from expansive self-renewal to differentiation. This altered fate is associated with reduced CSC proliferation in some but not all models. The data are consistent with a mathematical modeling study suggesting that negative feedback mechanisms in normal tissues may persist to some extent in tumors, with feedback on stem cells operating on both the probability of differentiation and the rate of self-renewal^18^.

### CSCs are hyper-responsive to microenvironmental inputs

Increasing culture density is associated with many microenvironmental changes including increased cell-cell contact and cell crowding, reduced cell spreading and cell-matrix interaction, depletion of nutrients and bioactive factors, and buildup of toxic metabolites or inhibitory factors. We first addressed the contribution of cell crowding. When LM2 CSCs in unsorted cultures reached their plateau, we removed the spent medium, trypsinized cells and reseeded into the same wells either at the original density (1-in-1) or at a 1-in-5 split ratio, in the presence of the original spent medium (**Fig. 4a**). The reduction in cell density from the 1-in-5 split led to a rapid expansion of the CSC population, while re-seeding at the original cell density had no effect. In contrast, the nonCSCs in the same culture showed little response to the low density reseeding. The data suggest that relief of cell crowding can selectively promote CSC expansive self-renewal. Cell crowding decreases ECM contact and reduces integrin-driven activation of FAK/SRC signaling ^19, 20^. The FAK inhibitor Defactinib slowed CSC population expansion (**Fig. 4b**), consistent with a positive role for cell spreading and integrin signaling in driving CSC expansive self-renewal.

**Fig. 4.**
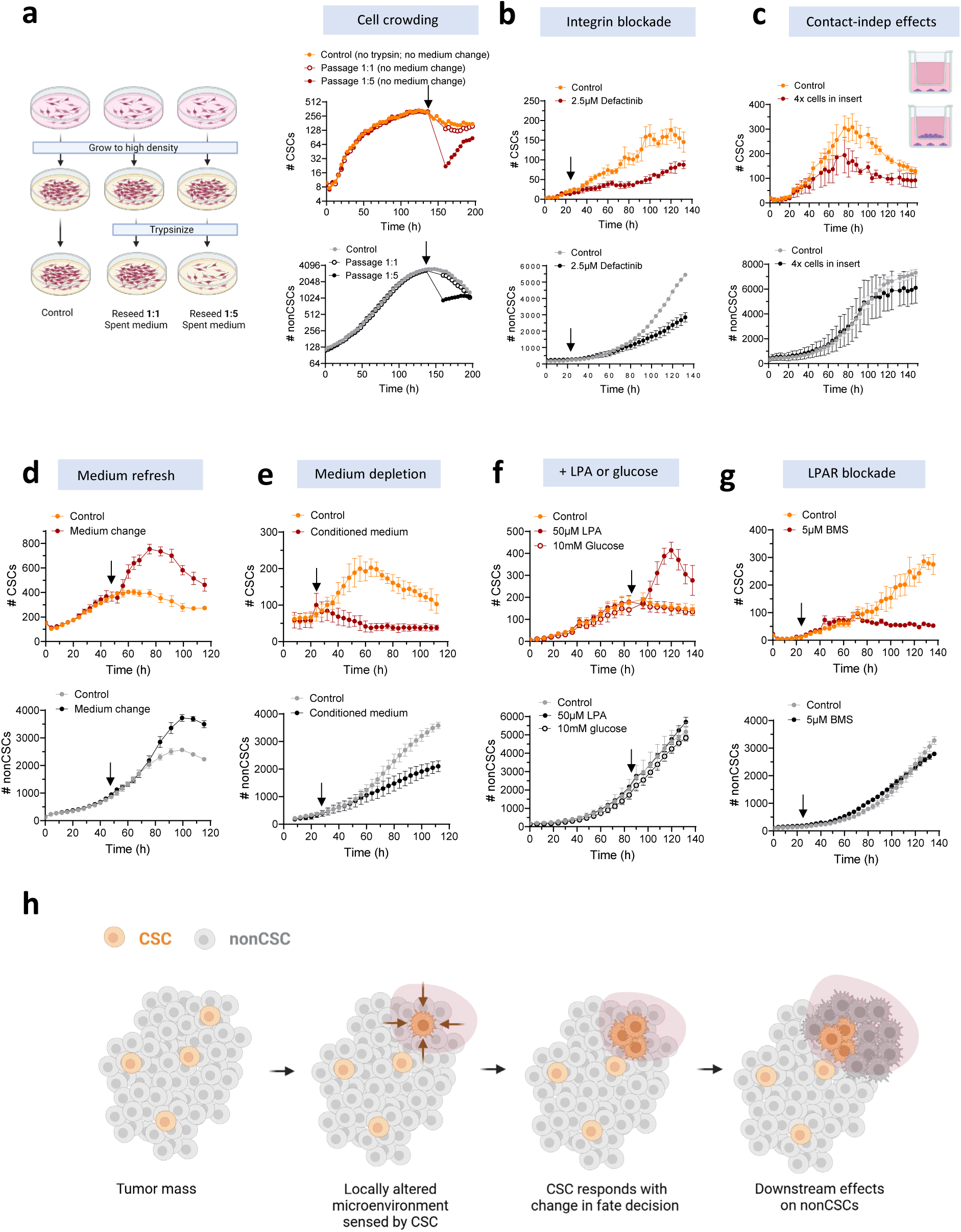
| CSCs are hyper-responsive to microenvironmental inputs. **a.** Experimental design and effect of relief of cell crowding on CSC (orange/red) or nonCSC (gray/black) population dynamics in unsorted LM2 cultures. Arrow indicates time of reseeding. **b.** Effect of integrin signaling blockade with FAK inhibitor (defactinib) on LM2 CSC and nonCSC dynamics. Arrow indicates time of inhibitor addition. **c.** Contact-independent effects on LM2 CSC and nonCSC expansion of additional high density LM2 cells in culture well insert. **d.** Effect on LM2 cells of replacing spent medium with fresh medium at 48h (arrow). **e.** Effect of switching medium at t=24h to medium pre-conditioned by high density cultures. **f.** Addition of 50µM LPA or 10mM glucose at 90h (arrow) to LM2 cultures. **g.** Addition of 5µM LPAR inhibitor BMS986020 (BMS) at 24h (arrow) to LM2 cultures. (**a-g**) Results are mean +/− SD (n=3). Results for CSCs and nonCSCs in the mixed cultures are plotted separately for clarity. **h.** Model for role of CSC as a microenvironmental sensor for the tumor cell culture/tumor mass.

Effects of culture density that were independent of cell crowding/spreading were also observed. The presence of tumor cells in a transwell insert decreased peak CSC numbers in the well below, while having relatively little effect on nonCSCs in the same culture (**Fig. 4c**). The contact-independent inhibitory effect could be due to build-up of inhibitory factors or depletion of stimulatory ones. Refreshing the medium when LM2 CSCs reached their plateau led to a transient increase in expansive CSC self-renewal, while having a smaller and more delayed effect on nonCSCs (**Fig. 4d**). Conversely, treatment of expansion phase CSCs with conditioned medium from high density cultures led to CSC loss (**Fig. 4e).** The stimulatory effect of medium refreshment on CSCs could be mimicked by addition of lysophosphatidic acid (LPA), a major serum mitogen^21^, but not by additional glucose (**Fig. 4f**). Conversely, CSC expansion in low density culture was blocked by the LPA receptor inhibitor BMS986020 (**Fig. 4g**).

Taken together, the results suggest that CSCs are highly sensitive to density-dependent changes in cell-matrix interaction, cell crowding and medium composition. In every case we noted that the response of CSCs to a perturbation was more extensive and/or faster than the response of nonCSCs (**Fig. 4b-g**), suggesting that CSCs are hyper-responsive to microenvironmental change. Similar results were seen with 4T1 or SUM159 cells (shown for FAK inhibition and medium replenishment in **Extended Data Fig. 4a-d**). These observations led us to propose that the CSCs serve as sensitive sensors of microenvironmental quality for the tumor cell population as a whole **(Fig. 4h)**. In this scenario, the CSCs continuously evaluate their local microenvironment in different regions of the tumor. They rapidly respond with a decision about whether the local conditions will support tumor growth, and then calibrate their numbers in response to available resources and space by altering the balance between self-renewing and differentiating fate decisions. Ultimately since nonCSCs are derived from CSCs, these rapid CSC responses will read out in longer-term changes in the nonCSC population. However, this will occur with a temporal lag, as most nonCSCs have some residual proliferative capacity and nonCSC proliferation is less immediately affected by environmental conditions.

### Multiple CSC sensor inputs converge on YAP/TAZ

To identify potential molecular mechanisms mediating the enhanced CSC responses to the microenvironment, we performed RNA-Seq on LM2 CSCs and nonCSCs sorted from *in vitro* cultures or metastasis-bearing lungs. As expected, the CSC transcriptome *in vitro* and *in vivo* showed strong enrichment for gene signatures related to stemness, drug resistance and metastasis (**Fig. 5A and Extended Data Fig. 5a**). Geneset Enrichment Analysis (GSEA) identified the Hippo pathway as the most highly enriched Reactome Signaling pathway in the CSCs *in vitro* (**Fig. 5b**), and Ingenuity Pathway Analysis® identified the Hippo pathway effector YAP as the most significant positive upstream regulator of the CSC transcriptome (**Fig. 5c**). YAP and its close relative TAZ (WWTR1 gene) function as a key nodal point in the transduction of biomechanical and other extracellular cues that regulate organ size and cell fate decisions^20, 22, 23^. They have been previously implicated in regulating the stem phenotype in breast and other cancers^20, 22^, and are plausible integrators of inputs to the CSC in its role as a microenvironment sensor. The Hippo pathway is a key negative regulator of YAP/TAZ^24^. However, a recent phenotypic screen for small molecule inhibitors of YAP/TAZ activation identified blockers of Rho-GTPase signaling as top hits, highlighting Rho-GTPases as important positive upstream regulators of YAP/TAZ activity^25^. Both YAP and Rho-GTPase activity were preferentially enriched in the transcriptome of CSCs sorted from lung metastases *in vivo* (**Fig. 5d**), as well as *in vitro* (**Fig. 5b**).

**Fig. 5.**
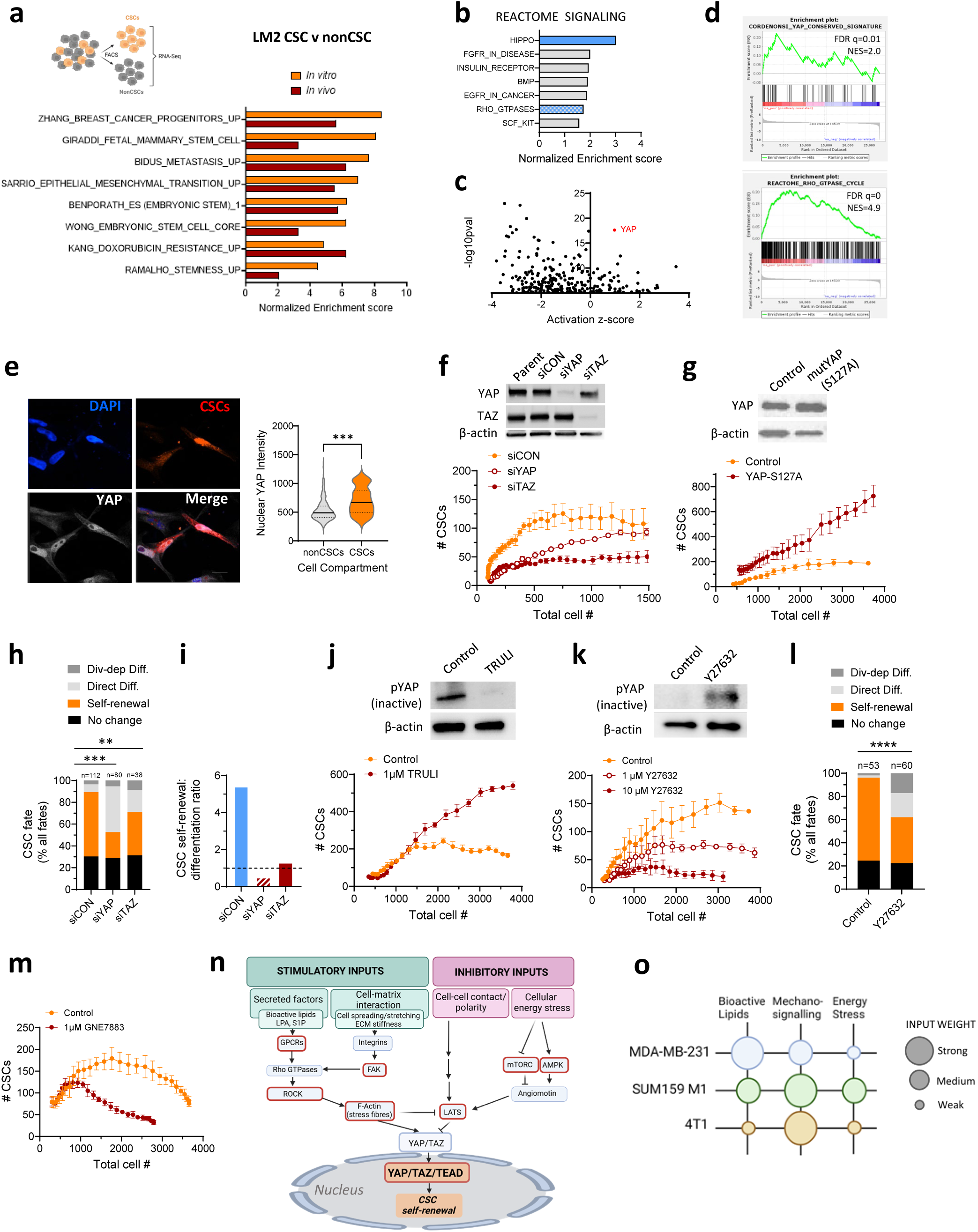
| Multiple microenvironmental inputs to the CSC sensor converge on YAP/TAZ. **a.** Enrichment of genesets relating to stemness in the transcriptomes of LM2 CSCs compared with nonCSCs, prepared from FACS-sorted tumor cells from *in vitro* cultures or recovered from metastatic lungs. **b.** Top Reactome Signaling pathways enriched in the CSC vs nonCSC transcriptome from sorted LM2 cells *in vitro.* **c.** Ingenuity Pathway Analysis showing predicted upstream regulators of the LM2 CSC transcriptome *in vitro*. **d.** Geneset enrichment plots for YAP signature and Rho-GTPase signaling in CSCs recovered from LM2 metastases *in vivo*. **e.** Quantitation of nuclear YAP intensity in CSCs and nonCSCs in unsorted LM2 cells in low density culture, using multiplexed immunofluorescence. n=24 cells (CSCs) and n=493 cells (nonCSCs); Mann-Whitney test. **f.** Effect of siRNA knockdown of YAP or TAZ on CSC dynamics in LM2 cultures. Note results are plotted on cell density axis (total number of cells). Results are mean +/− SD (n=3). **g.** Effect of overexpression of constitutively active mutant YAP-S127A on LM2 CSC dynamics. Results are mean +/− SD (n=3). **h.** Single cell fate mapping of LM2 CSCs following YAP or TAZ knockdown. Fisher Exact test vs control for self-renewal vs other fates. n = number of cells assessed. **i.** Effect of YAP/TAZ knockdown on CSC self-renewal:differentiation ratio in LM2 cells (data from **h**). **j,k.** Effect of LATS kinase inhibitor TRULI (**j**) or ROCK inhibitor Y27632 (**k**) on CSC dynamics in unsorted LM2 cultures. Results are mean +/− SD, n=3. Western blots show effect of inhibitors on phopho-YAP-S127 (inactive form of YAP). **l.** Single cell fate mapping showing effect of ROCK inhibition on CSC fate in LM2 cultures. Fisher Exact test for self-renewal vs other fates. n = number of cells assessed. **m.** Effect of TEAD inhibitor GNE7883 on CSC dynamics in unsorted LM2 cultures. Results are mean +/− SD, n=3. **n.** Schematic of upstream inputs to YAP/TAZ node. Red outlines indicate pathway components targeted by inhibitors in this study. **o**. Bubble plot summarizing relative weight of different input pathways across 3 models of TNBC, as assessed by response to inhibitors (see Suppl. Table 1 for detailed results).

Both the Hippo pathway and Rho-GTPases regulate YAP and TAZ through effects on their nuclear vs cytoplasmic localization and degradation^23^. Consistent with expectations of higher YAP/TAZ activity at low cell density, YAP/TAZ were preferentially localized in the nucleus of LM2 tumor cells in low vs high density cultures *in vitro* (**Extended Data Fig. 5b**), and in small vs large metastases *in vivo* (**Extended Data Fig. 5c**). Western blots confirmed increased phosphorylated (inactive) YAP in high density LM2 cultures, and in low density cultures treated with conditioned medium from high density cultures (**Extended Data Fig. 5d**). Importantly, immunofluorescent staining showed more nuclear YAP in CSCs than nonCSCs in LM2 cultures *in vitro*, indicating that YAP is preferentially activated in CSCs, consistent with the transcriptomic results (**Fig. 5e**).

To address their functional role, we showed that knockdown of YAP or TAZ in LM2 cells decreased CSC population expansion (**Fig. 5f**), while overexpression of the constitutively active mutant YAP-S127A increased CSC numbers and over-rode the density inhibition (**Fig. 5g**). Since YAP and TAZ can affect cell proliferation independently of their effects on cell fate decisions^22^, the data are plotted against total cell number (cell density axis) rather than time. This approach deconvolutes direct effects on CSCs from indirect effects that are secondary to changes in proliferation/density of the culture as a whole. Fate mapping showed that YAP or TAZ knockdown reduced CSCs by decreasing expansive self-renewal and increasing differentiation (**Fig. 5h,i**).

The Hippo pathway negatively regulates YAP/TAZ activity through activation of LATS kinases at high cell density^23^, while Rho-GTPases positively regulate YAP/TAZ through the effector kinase ROCK. ROCK plays a key role in mechanotransduction through effects on F-actin^24^, mostly independent of LATS^20^. YAP/TAZ then interact with TEAD family transcription factors to modulate transcription^26^. The LATS inhibitor TRULI blocked YAP phosphorylation and enabled continued expansion of CSCs at high density (**Fig. 5j**). Conversely, the ROCK inhibitor Y27632 inactivated YAP and reduced CSC expansion (**Fig. 5k, Extended Data Fig. 5e**), by increasing CSC differentiation and reducing self-renewal (**Fig. 5l**). Finally, pharmacologic blockade with the TEAD inhibitor GNE7883^27^ reduced CSCs (**Fig. 5m).** Thus both Hippo signaling and Rho GTPase signaling were confirmed as upstream regulators of CSC dynamics by modulating YAP/TAZ/TEAD activity.

Multiple stimulatory and inhibitory inputs converge on YAP/TAZ ^20, 22, 23^ (schematized in **Fig. 5n**). To assess their relative importance, we tested inhibitors of several key nodes (red boxes in **Fig. 5n**) on CSC dynamics across three different models of metastatic TNBC. The results are summarized in **Extended Data Table 1**, and in the bubble plot in **Fig. 5o**. Essentially all YAP/TAZ inputs had some influence on CSC self-renewal, but the dominant input varied between models. For example, LM2 cells were particularly sensitive to depletion of bioactive lipids, possibly because the LPA synthesizing enzymes autotaxin (ENPP2) and phospholipase A1 (PLA1A) ^21^ are not expressed in LM2 cells (**Extended Data Fig. 5f**). In contrast, 4T1 CSCs are relatively insensitive to LPA (**Extended Data Fig. 5g,h;** compare with **Fig. 4g,h**). Overall our data suggest that the YAP/TAZ node in CSCs integrates information from multiple microenvironmental inputs, with relative input weight varying between models.

### CSC hyperresponsiveness involves amplified input signals and a more TEAD-accessible chromatin organization

The greater responsiveness of CSCs to microenvironmental signals could involve more sensitive detection of incoming signals leading to more active YAP/TAZ (input arm), or a more efficient conversion of YAP/TAZ activation into the phenotypic readout (effector arm). Using LPA as a representative stimulus in the LM2 model, we showed that CSCs have higher expression of the LPA receptor1 (LPAR1) on the cell surface (**Fig. 6a,b**), and LPA treatment increased nuclear YAP/TAZ to a greater extent in CSCs than nonCSCs (**Fig. 6c**). Thus LPA in the microenvironment generates an amplified signal in CSCs compared with nonCSCs.

**Fig. 6.**
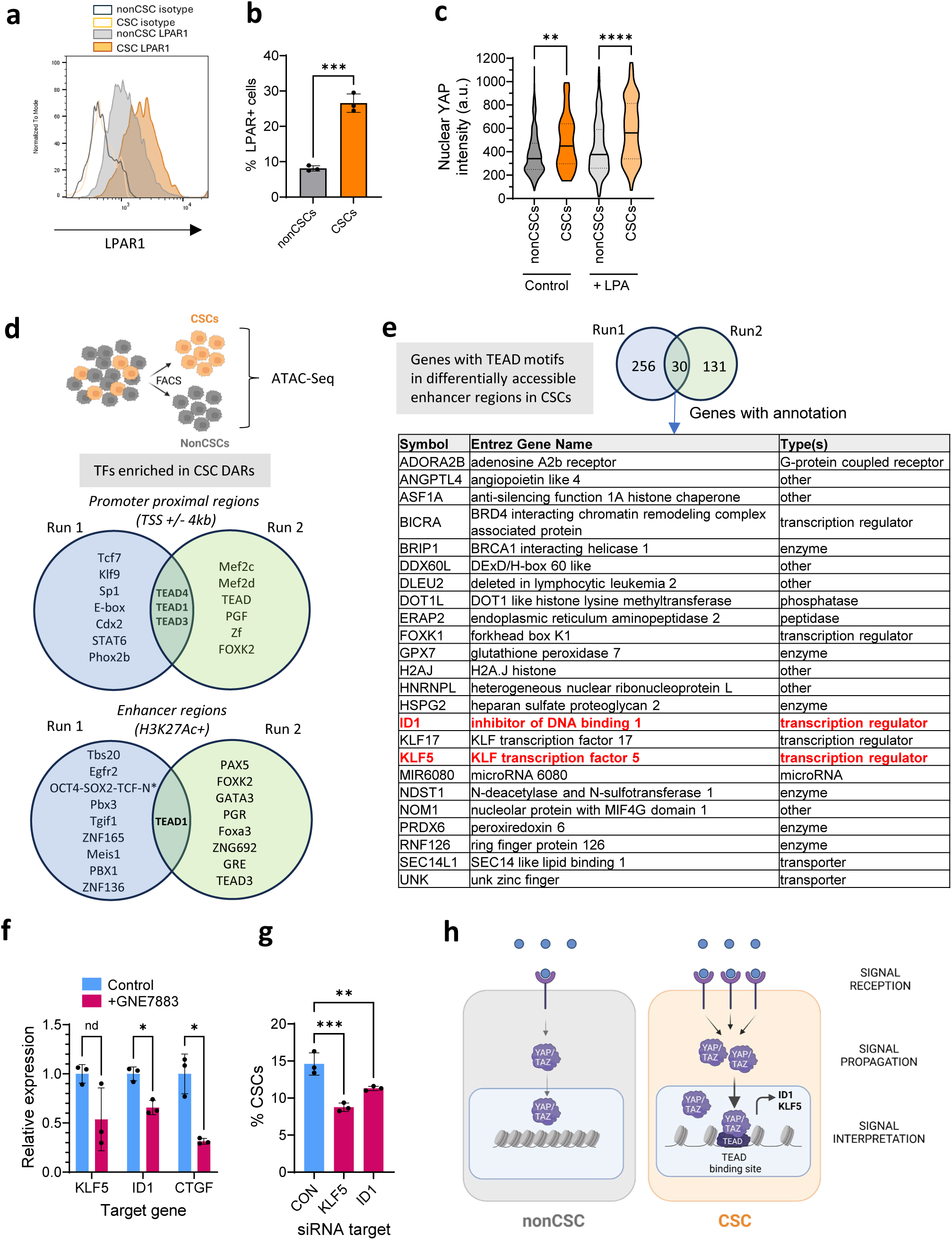
| CSC hyper-responsiveness to LPA involves amplified signaling and more TEAD-accessible chromatin. **a.** Flow cytometry for LPAR1 expression in LM2 CSCs (mCherry^pos^) vs nonCSCs (mCherry^neg^). **b.** Quantitation of data in (a). Mean +/− SD, n=3 **c.** Quantitation of YAP localization in LM2 CSCs vs nonCSCs in unsorted cultures with and without treatment with LPA assessed by immunofluorescent staining (n=524 and 400 for nonCSCs −/+ LPA; n=55 and 80 for CSCs −/+ LPA). Kruskal-Wallis test for CSCs v nonCSCs within treatment group. **d.** Top 10 enriched transcription factor binding motifs in differentially accessible chromatin regions in sorted LM2 CSCs vs nonCSCs around transcriptional start sites (TSS) or enhancer regions for 2 independent ATAC-Seq experiments. N* = NANOG. (Run 2 only had 9 enriched motifs that were statistically significant.) **e.** Genes with TEAD binding sites in their enhancers that are preferentially accessible in CSCs. Table shows overlap results of two independent runs (annotated genes only). **f.** RTQ-PCR for effect of TEAD inhibitor GNE7883 treatment on expression of KLF5 and ID1 in LM2 cells. CTGF is a canonical YAP target as positive control. Results are mean +/− SD for n=3, normalized to vehicle control. Students t-test. **g.** Flow cytometry for effect of 1D1 or KLF5 knockdown on % CSCs in LM2 cultures. Results are mean +/− SD for n=3, Dunnets multiple comparisons test vs control (CON). **h.** Schematic for mechanisms underlying hypersensitivity of CSCs to microenvironmental signals.

To address possible differences in the effector arm, we performed ATAC-Seq on sorted LM2 CSCs and nonCSCs. We did not see consistent global chromatin accessibility differences between the two subpopulations, but each had some local differentially accessible regions (DARs). Focusing on CSC DARs in promoter proximal regions (TSS +/− 4kb) and in previously identified MDAMB231 enhancers^28^, we searched for enriched transcription factor binding motifs. TEAD motifs were consistently enriched in CSC DARs at promoters and enhancers in each of two independent experiments, while other motifs were more variable (**Fig. 6d**). Since >90% of YAP/TAZ/TEAD binding occurs at distal active enhancer regions (marked by H3K27Ac) ^26, 29^, we focused on CSC enhancer DARs with TEAD binding sites. Among the genes associated with these enhancers (**Fig. 6e**), we found ID1, a well characterized inhibitor of differentiation^30,31^, and KLF5, a key transcription factor in the maintenance of the stem state^32^. Treatment of LM2 cultures with the TEAD inhibitor GNE-7883 reduced expression of ID1 and KLF5 expression (**Fig. 6f**), and knockdown of either ID1 or KLF5 with siRNA reduced CSCs (**Fig. 6g**), consistent with TEAD-dependent expression of both proteins maintaining stemness in LM2 cells.

Taken together, our data suggest that the ability of CSCs to act as early sensors of microenvironmental change is due to their greater sensitivity to incoming signals (enhanced signal reception and propagation), as well as an altered chromatin architecture that increases accessibility of YAP/TAZ/TEAD binding sites in regulatory elements of many target genes (enhanced signal interpretation) **(Fig. 6h)**, including those that inhibit differentiation and sustain stemness.

### Targeting CSC sensor nodes reduces chemotherapy-driven enrichment of CSCs in early metastasis

Many studies have shown that the CSC population is relatively resistant to therapy^8, 33^. Notably breast tumor tissue remaining after endocrine therapy or taxane chemotherapy was enriched for stem features^34^. Treatment of the LM2 model with Paclitaxel significantly reduced metastatic lesion size (**Fig. 7a,b**). However, Paclitaxel increased the proportion of CSCs among surviving cells (**Fig. 7c**), and this effect was seen across all lesion sizes (**Fig. 7d**). We therefore asked whether blocking CSC self-renewal could reverse this treatment-induced enrichment of CSCs. In principle, the YAP/TAZ/TEAD node would be the optimal target, and promising new pan-TEAD inhibitors have recently been developed^27^ ^35^. We tested GNE-7883 with Paclitaxel in the LM2 model and saw a significant decrease in metastasis number with the combination treatment (**Extended Data Fig. 7a,b**). However, the recommended drug excipient (sunflower oil^27^) interfered with our ability to image the CSCs reliably. Instead, for proof-of-concept we used Y27632 to inhibit ROCK, an upstream node for both bioactive lipid and mechano-signaling to YAP/TAZ (**Fig. 7e**). Flow cytometry *in vitro* confirmed that treatment with Paclitaxel alone selectively reduced nonCSCs in the LM2 model, while Y27632 alone selectively reduced CSCs, and the combination reduced both **(Fig. 7f)**. Extending these results to the *in vivo* setting, we showed that ROCK inhibition did not interfere with the ability of Paclitaxel to reduce metastatic lesion size (**Fig. 7g,h**) but did reverse the Paclitaxel-induced enrichment of CSCs across all lesion sizes (**Fig. 7i,j**). Combination therapy also increased the proportion of metastases that had no CSCs **(Fig. 7k)**, suggesting that some metastases were being “sterilized” of their CSCs. Similar results were seen with the LPAR inhibitor BMS5986020 in the LM2 model (**Extended Data Fig. 7c-g**). Thus therapeutic targeting of CSC sensor nodes combines with conventional chemotherapy *in vivo* to improve anti-metastatic responses.

**Fig. 7.**
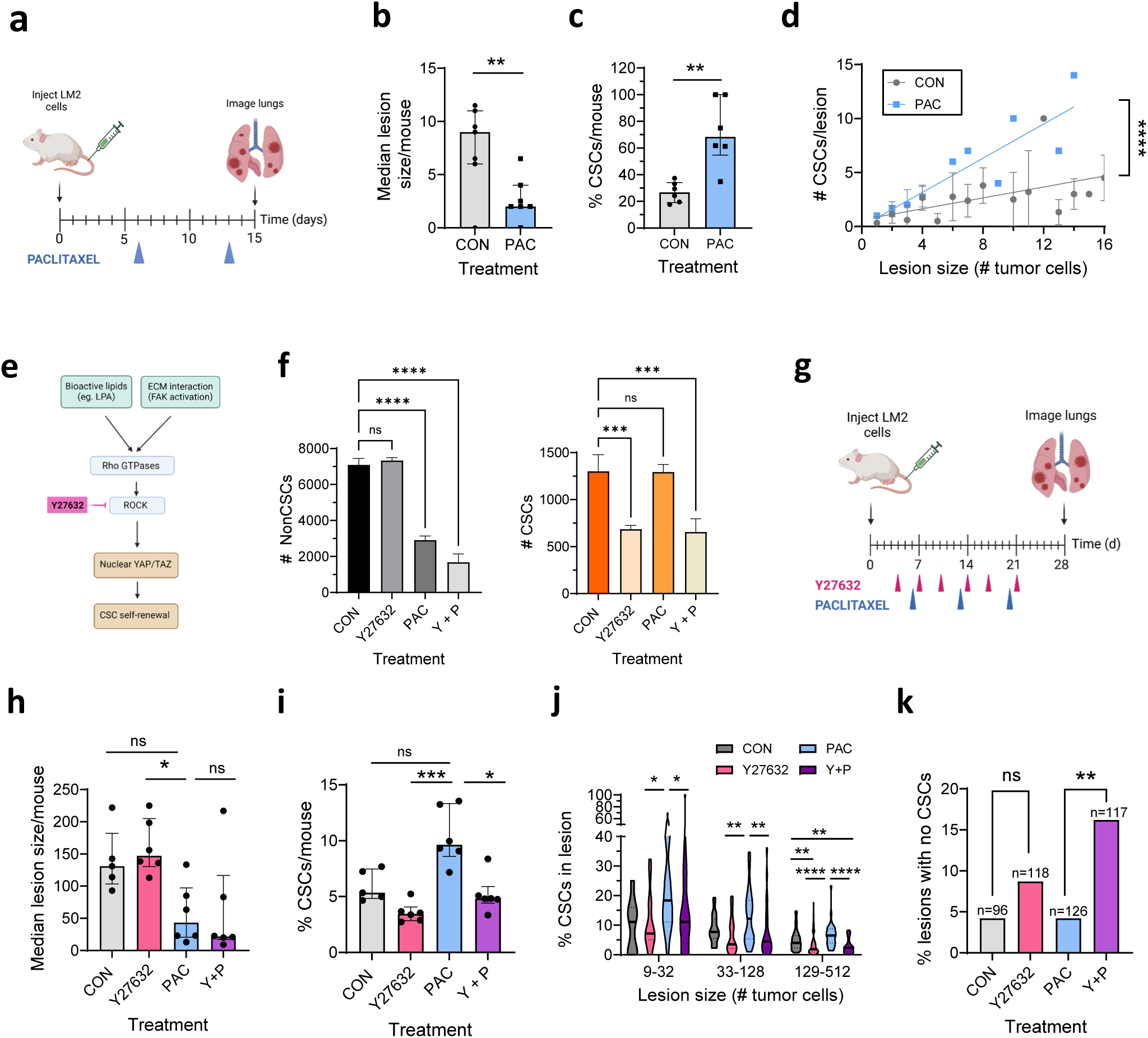
| Targeting CSC sensor nodes reduces chemotherapy-driven enrichment of CSCs in metastasis. **a.** Schematic for Paclitaxel treatment. **b-c.** Effect of Paclitaxel on LM2 lung metastasis size **(b)** and % CSCs among tumor cells in the lung **(c)**. Results are median +/− IQ range, n=6-7; Mann-Whitney test **d.** Number of CSCs/lesion as a function of LM2 lung lesion size in response to Paclitaxel. Results are mean +/− SD for n=1-10 lesions/size bin. ANCOVA test. **e.** CSC sensor inputs converging on ROCK. **f.** Effect of combined Paclitaxel + ROCK inhibitor treatment for 3 days on CSCs and nonCSCs in unsorted LM2 cultures *in vitro* assessed by flow cytometry. Results are mean +/− SD for n=3, Dunnett’s multiple comparison test vs CON. **g**. Schematic for Paclitaxel + Y27632 treatment in LM2 model. **h,i**. Effect of therapeutic interventions on median lung lesion size (# tumor cells)/mouse (**h**) or % CSCs/lung (**i**) at endpoint. Results are median +/− IQ range for n=5-6 mice/group; Dunn’s multiple comparisons test vs Paclitaxel. **j.** Treatment effect on % CSCS as a function of lesion size. Data pooled from 5-6 mice/group, n=8-70 lesions/size bin; Dunn’s multiple comparisons test. **k.** Treatment effect on % lesions with no CSCs. Fisher’s Exact test for indicated treatment pairs. n= total lesion number as indicated.

## DISCUSSION

### CSC dynamics in lung colonization

Direct observation of CSCs in metastatic breast cancer models using our fluorescent CSC reporter has established a detailed ground truth for CSC and nonCSC dynamics in early lung metastatic colonization. We initially confirmed that CSCs are strongly enriched among circulating tumor cells, have a selective survival advantage on arrival in the lung, and are uniquely capable of initiating metastases, as has been seen previously by us and others^14,17,36, 37^. Downstream of those early events, we discovered very different population dynamics for CSCs and nonCSCs in developing metastatic lesions. In the first few cell divisions, only CSCs divide, undergoing expansive self-renewal and also dividing asymmetrically to generate nonCSC progeny. However, the CSC population rapidly plateaus and further metastasis growth is driven by expansion of the nonCSC population. These divergent dynamics suggested that CSCs and nonCSCs respond to local microenvironmental cues differently, with the CSC population uniquely experiencing early negative feedback control as metastases grow larger.

We anticipated that this negative feedback would involve reduced CSC proliferation, since the relative quiescence of CSCs has been proposed as a contributing mechanism for their higher chemotherapy resistance^38^. Unexpectedly we found that CSCs in late-stage metastases from aggressive tumor models were just as proliferative as nonCSCs, despite the CSC population size remaining essentially static. However, this observation makes intuitive sense since nonCSCs, by definition, lack self-renewal capacity and thus can only undergo limited divisions before exhausting their proliferative potential. Consequently, ongoing CSC proliferation is required to replenish the proliferating nonCSC pool in expanding metastases. This outcome can be achieved without expansion of the CSC population if CSC self-renewal is balanced by CSC differentiation in larger lesions. Thus the negative feedback on CSCs in growing metastases primarily affects CSC fate decisions rather than their proliferation. The observation that CSCs in large metastases are at least as proliferative as nonCSCs importantly suggests that their relative resistance to anti-mitotic therapies relies on properties other than just quiescence.

### CSCs as first responders to a changing microenvironment

Modeling these distinctive population dynamics *in vitro*, we found that CSCs responded to many features of a crowded cell environment (eg. reduced matrix attachment, depleted trophic factors) with increased differentiation and reduced expansive self-renewal. Importantly, the CSC population consistently responded much earlier and more extensively than nonCSCs to a wide variety of stimuli. This observation led us to the broader concept that CSCs may serve as sensitive sensors and first responders to changing local microenvironmental conditions. Since CSCs are distributed through the tumor (see **Fig. 1c,d**), they are particularly well-situated to serve this role for the tumor as a whole. The ensuing CSC population expansion or contraction subsequently reads out in changes in the nonCSC progeny, and couples local tumor expansion dynamics to resource availability and biomechanical cues. This situation has clear analogies with developmental and repair processes, where mechanisms must be in place to integrate signals from the tissue as a whole, so as to calibrate tissue expansion to the functionally optimal size^23^. As an aside, it is worth noting that important experimental consequences of the high sensitivity of CSCs to microenvironmental inputs include (i) a strong dependence of CSC numbers and dynamics on initial conditions (“butterfly effect” in chaos theory), and (ii) culture density as a potentially confounding variable in many CSC assays and drug screens.

### The YAP/TAZ/TEAD node integrates microenvironmental inputs and generates amplified responses in CSCs

We found that microenvironmental inputs affecting breast CSC dynamics activated many interconnected upstream pathways that converged on the YAP/TAZ/TEAD node to alter the balance of self-renewal vs differentiation fate decisions in the CSCs. The Hippo pathway and its downstream effectors YAP,TAZ and TEADs are known to respond to biomechanical, metabolic, inflammatory and other cues to modulate cell fate decisions during development and tissue repair^23^. Earlier studies have implicated YAP and TAZ in regulating the stem phenotype in tissue regeneration and disease ^20, 22^. For example, YAP/TAZ reprogram adult intestinal cells into a fetal-like regenerative state in mouse models of intestinal damage^12^, and overexpression of TAZ conferred stem properties on oncogenically-initiated breast epithelial cells^39^. However, the selective hyper-responsiveness of this signaling node in CSCs that we identified in this study is a novel finding with important conceptual and practical implications for understanding CSC interactions with their changing microenvironment. Underlying mechanisms included enhanced signal reception and propagation in CSCs, and greater accessibility of TEAD binding sites in enhancers of stem cell regulatory genes such as KLF5 and ID1. Thus in comparison to nonCSCs, CSCs are primed to amplify responses to incoming microenvironmental signals at every level of an integrated sensor and effector network centered on YAP/TAZ/TEAD activity.

### Leveraging CSC response mechanisms to enhance therapeutic efficacy

Most current therapeutic approaches are relatively ineffective against CSCs, leading to CSC enrichment in post-treatment samples^1^. Conceptually, strategies to differentiate CSCs should prevent this phenomenon and lead to more durable responses. The paradigm for effective differentiation therapy is exemplified by the treatment of acute promyelocytic leukemia with retinoic acid and arsenic^40^. However, the mechanism in that setting probably involves degradation of the oncogenic fusion protein, and differentiation therapy has yet to prove efficacious in solid tumors with more complex genomes^40^. Furthermore, the critical goal must be CSC differentiation with loss of self-renewal capacity, rather than just differentiation of more committed progenitor cells, which will not lead to long-term cures. Pathways such as Wnt and Notch have previously been implicated in regulating CSC dynamics, but targeting these has proved challenging due to their involvement in normal tissues^41^. In contrast, the hypersensitivity of CSCs to microenvironmental signals that regulate CSC self-renewal via YAP/TAZ/TEAD presents an exploitable vulnerability, since YAP and TAZ are largely dispensable for normal adult homeostasis ^20, 22^.

Newly developed TEAD inhibitors are showing early promise against Hippo pathway-mutated tumors, and as a strategy to circumvent resistance to targeted therapies^27, 35^. While the effects of TEAD inhibition on CSCs were not directly assessed in those studies, the major role for CSCs in therapy resistance makes them plausibly the critical cellular targets. In our hands, the TEAD inhibitor GNE-7883 effectively reduced CSCs *in vitro* and showed some efficacy in a preclinical metastasis model, despite drug delivery challenges. In an alternative proof-of-principle experiment using a ROCK inhibitor to block upstream signaling to YAP/TAZ/TEAD, we effectively prevented the paclitaxel-driven enrichment of CSCs in lung metastasis models. Since we showed that the relative weight of upstream inputs to the YAP/TAZ/TEAD node varies among tumor models, direct inhibitors of YAP/TAZ/TEAD should be the most broadly applicable therapeutic strategy, if safety profiles are tolerable. However, the possibility of using a bet-hedging strategy by simultaneously targeting multiple input nodes upstream of YAP/TAZ is worth further investigation. A potential advantage of a low dose multitarget approach would be divided toxicity and reduced activation of bypass mechanisms, as was observed with a multidrug strategy to overcome variations in MAPK network topology^42^. Since the efficacy of any treatment that differentiates CSCs will be limited if the rate of de-differentiation of nonCSCs is appreciable, therapeutics that induce CSC differentiation and simultaneously block nonCSC plasticity would be optimal. Our dynamic reporter with its ability to report on both processes should help in the quest for such agents.

## Methods

### Cell lines and cell culture

The origin, providers and culture media for breast cancer cell lines used in this study were as follows: MDAMB231 (human triple negative breast cancer (TNBC); American Type Culture Collection; cultured in DMEM/10% FBS); MDAMB231-LM2^15^ (lung-tropic variant of parental MDAMB231; Dr. Joan Massague; DMEM/10% FBS); SUM159M1a^43^ (human anaplastic TNBC; Dr. Yibin Kang; Ham’s F-12/10% FBS); SUM149PT^44^ (human BRCA1-mutant inflammatory breast cancer; Dr. Esta Sterneck; Ham’s F-12/10%FBS, 5ug/ml insulin, 1ug/ml hydrocortisone); MCF10Ca1h^45^ (tumorigenic Ras-transformed human mammary epithelial cell; Dr. Fred Miller, DMEM/F12/5% horse serum; 4T1^16^ (mouse TNBC; Dr. Fred Miller, DMEM, 10% FBS). Cells were cultured in a humidified incubator at 37°C with 5% CO_2_. All human cell lines underwent authentication via short tandem repeat (STR) profiling (TransnetYX Inc, CA). Mycoplasma testing was performed regularly using the Lookout Mycoplasma PCR Detection Kit (Sigma-Aldrich, St.Louis MO).

### Pharmacologic inhibitors

The pharmacologic inhibitor source, dose range and vehicle used for *in vitro* studies were as follows: TRULI (Selleckchem, E1061, 1-10 µM, DMSO); Defactinib (Selleckchem, S7654, 2.5-12.5 µM, DMSO); Cytochalasin D (Selleckchem, S8184, 0.05-1 µM, DMSO); BMS986020 (Selleckchem, S3572, 1-10 µM, DMSO); AM966 (Selleckchem, S3534, 1-25 µM, DMSO); Y27632 (Selleckchem, 1-10 µM, DMSO); H1152 (Selleckchem, 0.5-5 µM, S6214, DMSO); Fasudil (Selleckchem, S1573, 1-50 µM, DMSO); Torkinib (Selleckchem, S2218, 50-500 nM, DMSO); AICAR (Selleckchem, S1802,0.1-10 µM, PBS); MK3903 (Selleckchem, S8803, 0.5-10 µM, DMSO); GNE7883 (MedChem Express, HY-147214, 0.5-5 µM, DMSO); Paclitaxel (Selleckchem, S1150, 1-25 nM, DMSO)

### Reporter constructs and lentiviral transduction

All cell lines were transduced with a constitutively expressed nuclear GFP marker using EF1alpha>nucGFP lentivirus (Cat #4624 Essen Bioscience, CA) and selected by FACS. The SORE6>dsmCherry pro-lentiviral reporter construct (18065-M02-412) and minCMV>dsmCherry control construct (18065-M04-412)) were modifications of our previously described constructs ^13, 14^, using a destabilized mCherry in place of dsCopGFP, (**Extended Data Fig. 1**). Titered lentivirus was generated by Applied StemCell Inc, CA. Exponentially growing cells already expressing the constitutive nucGFP were transduced by exposure to virus at an MOI of 1 for 24h without polybrene, and then expanded and FACS sorted for CD19 expression to ensure integration of reporter lentivirus in all cells. The low MOI is critical to restrict reporter construct integration to one copy/cell. Cells were typically used within 10 passages from reporter transduction.

### Animal models and metastasis assays

All animal experiments were conducted in compliance with institutional guidelines and under protocol LC-070, approved by the Institutional Animal Care and Use Committee (IACUC) of the Center for Cancer Research (CCR), National Cancer Institute. The CCR is accredited by AALAC International and follows the Public Health Service policy for the Care and Use of Laboratory Animals. Virgin female NCI Ath/Nu++ immunodeficient nude mice and immunocompetent BALB/c mice were obtained from NCI Charles River Labs at 6-8 weeks of age. All experiments involving cells containing imaging markers were performed in the immunodeficient mice to avoid rejection of the fluorescent protein neoantigen. Several different metastasis assay formats were used. For MDAMB231-LM2 cells, the orthotopic format involved orthotopic implantation of 5 x 10^5 labelled cells in 0.1 ml of PBS by injection into the #2 and #7 mammary fat pads. For the pseudo-orthotopic format, 2.5×10^5 unlabelled cells were injected into the #2 mammary fat pad to establish a pre-metastatic niche and then after 10-15 days, 4 x 10^5 labeled cells were injected via the tail vein in 100ul of sterile phosphate-buffered saline (PBS). Lungs were harvested at various timepoints after tail vein injection. For the 4T1 model, 4 x 10^4 labeled cells were surgically implanted in the #2 and #7 mammary fat pads of nude mice. After 14 days, primary tumors were resected. Lungs were harvested at various timepoints after primary tumor resection.

### *In vivo* drug treatment studies

The TEAD inhibitor GNE-7883 (Chemieteck) was suspended for *in vivo* use in sunflower oil (Spectrum Chemical Corp) and dosed daily by subcutaneous injection on the nape of the neck at 250mg/Kg for 4 days on, 2 days off according to Hagenbeek et al.^27^. The ROCK inhibitor (SelleckChem, Cat # S1049)) was prepared in saline and dosed at 10 mg/Kg i.p. twice/week^46^. Paclitaxel was obtained from the Division of Veterinary Resources at NCI. The 6mg/ml stock in Cremaphor was diluted to 2.8mg/ml in PBS and administered by intravenously via tail vein injection at 10 mg/Kg once/week. The LPAR inhibitor BMS5986020 (SelleckChem Cat#S3572) was prepared in 0.5% methylcellulose in water and delivered by oral gavage daily^47^.

### Circulating tumor cells

Mice were orthotopically implanted with LM2 or 4T1 cells transduced with the nuclear GFP constitutive marker and the SORE6>dsmCherry CSC reporter. When tumors reached 0.5-1.0 cm diameter, mice were anesthetized and 1ml of fresh blood was collected via cardiac puncture into EDTA (1.6mg/ml) tubes. Samples were then processed using the Cytiva Ficoll-Paque PLUS (GE Healthcare, Chicago, IL, USA) cell preparation method according to manufacturer’s instructions. 0.1ml of the buffy coat suspension was centrifuged at 300rpm for 5 mins using a Cytospin cytocentrifuge to deposit the cells onto glass coverslips. After brief air-drying, the slides were scanned using an EVOS FL Auto 2 Imaging System (Thermo Fisher Scientific, Waltham, MA, USA) to identify and quantify GFP-positive cancer cells and mCherry-positive cancer stem cells (CSCs). Higher resolution images were acquired using Nikon SoRa spinning disk confocal microscope with 20x objective lens.

### Flow Cytometry and Fluorescence-Activated Cell Sorting (FACS)

#### Flow cytometry

Subconfluent cultures of cells were harvested by trypsinization for antibody staining and analyzed on a BD LSRFortessa SORP1 instrument (Becton Dickinson). Data acquisition and analysis were performed using FlowJo Software (Tree Star). Cell viability was assessed using 7-AAD staining. Cells transduced with minCMV>dsmCherry were used as the gating control for the SORE6>dsmCherry reporter. Antibodies used were: LPAR1 (Alomone Labs LTR, Cat# ALR-031).

#### FACS

SORE6-mCherry transduced cells were sorted using a BD FACSAria Fusion system. After gating to exclude doublets, dead cells (7-AAD staining) and CD19-cells, gating for SORE6>dsmCherry positivity was established using mCMV-dsmCherry-expressing cells as negative controls. The top 5% of cells in the SORE6+ gate and bottom 20% in the SORE6-gate were collected for further analysis or injection into mice. Cells recovered from primary tumors or metastatic lungs, were stained with CD19 antibody (APC-anti-human-CD19, Biolegend Cat#392504). Tumor cells were identified and gated based on GFP and CD19 positivity. CSCs and nonCSCs were further distinguished using the mCherry fluorescence gate.

### Incuctye Live Cell Imaging

#### CSC and nonCSC dynamics

An Incucyte S3 Live-Cell Imaging system (Essen Biosciences) with 440/480 ex/em (green) and 565/620 ex/em (red) filters was used for live-cell imaging to monitor CSC and nonCSC dynamics in culture over time. Cells were plated in standard 24-well tissue culture plates (Corning Cat# 3524). Images were captured in the red and green channels every 4 h for the duration of each experiment. Images were analyzed using the basic analyzer in the Incucyte 2023 Rev B software (Sartorius). Cells expressing mCherry above a predefined threshold were classified as CSCs. This threshold was determined using the minCMV>dsmCherry control, set to ensure <0.2% of total cells were red-positive in the control sample. Cells only expressing GFP and not mCherry were considered nonCSCs. In many experiments, analysis definitions were validated by flow cytometry data confirming correct % CSCs at the endpoint.

#### Live Cell Fate-mapping

For CSC fate and origin mapping, LM2 cultures were imaged in the Incucyte S3 system as above, except that imaging occurred every 30 mins, starting at 6h after plating at 25,000 or 50,000 cells/well in 24-well plates in DMEM/10% FBS. Using cell analysis definitions established as above, SORE6+ cells were masked in red on the images at each timepoint. Movies were generated to allow tracking of cells that were SORE6+ (CSCs) at the start of the imaging period to determine their fate over the following 48-72h. Possible fates are schematized in **Fig. 3l**. SORE6+ cells were only considered to have differentiated if they maintained loss of red signal for ≥3 hours of tracking (6 consecutive frames). Fate decisions of SORE6+ cells were assessed by mapping 20-40 SORE6+ cells/well from 3-4 wells/condition, resulting in a total of 60-110 cells/treatment group. Origin mapping of SORE6+ CSCs was performed similarly, but starting with SORE6+ cells at the end of the mapping period and tracing back to their origin at earlier times.

### siRNA-mediated knockdown

siRNA-mediated knockdown was performed using SMARTpool ON-TARGETplus siRNAs (Horizon Discovery) targeting YAP (catalog # L-012200), TAZ (L-016083), LATS1 (L-004632), KLF5 (L-013571), and ID1 (L-005051). MDA-MB-23LM2 cells were transfected with siRNAs using Lipofectamine RNAiMAX (Thermo Fisher Scientific) according to the manufacturer’s instructions. Briefly, cells were seeded in antibiotic-free medium 24 hours prior to transfection. siRNA-lipid complexes were formed by mixing siRNAs with Lipofectamine RNAiMAX in Opti-MEM medium (Thermo Fisher Scientific) and incubated for 20 minutes at room temperature before adding to cells. Knockdown efficiency was assessed by Western blot 72 hours post-transfection.

### RTQ-PCR

Total RNA was isolated using the RNeasy Mini Kit (QIAGEN) according to the manufacturer’s instructions. Reverse transcription was performed using the SuperScript III RT kit (Thermo Fisher Scientific) to generate cDNA. Quantitative PCR was conducted using the SoFast qPCR Mastermix (Bio-Rad) on a real-time PCR system (Bio-Rad CFX Opus96). All reactions were performed in triplicate, and relative gene expression was calculated using the 2^(-ΔΔCt) method. PP1A was used as the reference gene for normalization. Primers were purchased from IDT, and the sequences are given in Supplementary Information.

### RNA sequencing

#### Isolation of RNA

RNA was prepared from cell cultures using the Trizol Reagent (Ambion) according to manufacturer’s instructions. For RNA isolation from sorted cells, cells were sorted directly into Trizol. RNA quality was assessed using the Agilent 2100 Bioanalyzer system. All RINS were 9 or above. For RNA preparation from cells in metastatic lungs, tissues were dissociated into single-cell suspensions by mechanical processing through 70 μm and 40 μm filters, followed by centrifugation. Cells were resuspended in PBS with 4% FBS, stained with CD19-APC, and sorted via FACS based on GFP+, CD19-APC+, and mCherry gates. Sorted cells were collected directly into Trizol for RNA isolation.

#### Sequencing

RNA-Seq was performed at the Next Generation Sequencing Facility, Center for Cancer Research at the NCI. Sequencing libraries were generated using Illumina TruSeq® Stranded Total RNA Kit RS-122-2201 protocol and run on the Illumina HiSeq platform with HiSeq 3000/4000 sequencing chemistry.

#### Bioinformatic analysis

Sample reads were trimmed for adapters and low-quality bases using Cutadapt before alignment with the human reference genome (hg19). Differential gene expression was performed using DESeq2 to compare CSC vs. nonCSC for the LM2 cell line.

#### Pathway and Geneset Enrichment analysis

Differentially expressed genes with an FDR cut-off of 0.25 were analysed using Ingenuity Pathway Analysis Software (Qiagen) to determine the top predicted Upstream Regulators of the CSC and nonCSC genesets. Gene set enrichment analysis was performed using GSEA based on the entire gene list ranked based on the statistics from the DESeq2 analysis in which we added the fetal mammary stem cell signature^48^ to the curated gene sets (C2).

### ATAC-Seq

#### ATAC-Seq Library Preparation and sequencing

Samples were processed using the Active Motif ATAC-Seq Kit (Active Motif Inc, Carlsbad CA; #53150) according to the manufacturer’s instructions. Two independent preparations of FACS-sorted LM2 SORE6+ and SORE6-cells that had been allowed to recover in culture overnight (four samples total) were used. Briefly, 100,000 cells were collected and resuspended in 100 µl of ice-cold ATAC Lysis Buffer. The cell suspension was centrifuged at 500 × g for 10 minutes at 4°C. The resulting cell pellet was resuspended in 50 µl of Tagmentation Master Mix and incubated at 37°C for 30 minutes. The tagmented DNA mixture was combined with 250 µl of DNA Purification Binding Buffer and 5 µl of 3 M sodium acetate, followed by purification using a DNA purification column as per the manufacturer’s instructions. The purified DNA was then amplified using indexed primers via PCR and sequenced on a NovaSeq 6000 SP.

#### ATAC-Seq analysis

All analyses were carried out on NIH Biowulf2 high performance computing environment using software default parameters, unless specified otherwise. The fastq sequence reads were aligned to the reference genome hg38 using Bowtie (2-2.5.3) mapping software. Peak calling was performed using MACS2 (2.2.7.1) software with the parameters of –broad –nomodel. Heatmaps were generated using deepTools (3.5.5). Analysis used bamCoverage, computeMatrix, and plotHeatmap from deepTools. Transcription factor binding sites were analyzed with Homer (5.1). findMotifsGenome.pl was used to find transcription factor motifs that are enriched in the ATAC-seq peaks for CSC vs. nonCSC, and scanMotifGenomeWide.pl was used to map transcription factor binding sites for the motifs that are enriched in CSC. H3K27Ac ChIPseq data for MDAMB231 cell line was downloaded from SRA database (SRR949140). It was aligned with Bowtie with hg38, and peak calling was performed using MACS2. The peak regions were used to define enhancer regions in the LM2 subline.

### Tissue preparation for imaging and immunostaining

#### Primary tumors

Upon reaching the desired size (∼5mm diameter) tumors were surgically excised and immediately placed in ice-cold phosphate-buffered saline (PBS). Tumor tissues were then cut into 1-2mm slices with a razorblade and mounted onto glass-bottom dishes for confocal imaging.

#### Lung metastases

Following euthanasia by CO2 asphyxiation, the lungs were inflated with ice-cold PBS and then carefully excised and the same one lobe from each lung was selected for fresh tissue imaging. This lobe was gently laid flat onto a glass-bottom Ibidi 35 mm dish (Ibidi GmbH, Martinsried, Germany), minimizing handling to prevent deformation and immediately imaged by confocal microscopy. The remaining lobes were either fixed in formalin for histologic analysis, or embedded in OCT compound (Tissue-Tek, Sakura Finetek USA Inc, Torrance, CA) and frozen for immunofluorescent staining.

### Confocal imaging and analysis of tissues

#### Tissue imaging

Confocal imaging of lung metastases was mostly performed using a Zeiss LSM 880 confocal microscope (Carl Zeiss Microscopy GmbH, Jena, Germany) equipped with a GaAsP spectral detector and plan-apochromat 40x/1.3 oil DIC UV-IR M27 objective lens for enhanced sensitivity and high resolution confocal images. 25-50 random fields of view per lung were scanned and images were captured for analysis. Data were normalized to the total number of fields scanned. For imaging primary tumors and metastatic lungs over a wider area, a Nikon SoRa spinning disk confocal microscope equipped with photometrics BSI sCMOS camera and 20x plan-apochromat (N.A. 0.75) objective lens was used in tile imaging mode. Image tiles were merged using the stitching algorithm in the Nikon NIH-Elements software (v.5.4.2). The microscope setup allowed simultaneous excitation of GFP and mCherry, using 488 nm and 561 nm lasers, respectively. Images were acquired with NIS-Elements software (Nikon Corporation, Tokyo, Japan) and processed to enhance clarity and contrast using Imaris software (Oxford instrument, Oxfordshire, England)

#### Tissue image analysis

Captured images were processed and analyzed using ImageJ software to quantify metastatic lesion size and CSC numbers. Metastatic lesions were identified based on the fluorescence intensity of GFP, which marked the nuclei of all cancer cells. Using ImageJ Fiji the total number of cancer cells in each individual lesion was counted. The number of CSCs (SORE6>dsmCherry positive) in each lesion were assessed using mCherry signal segmentation after setting an mCherry threshold on lesions from tumor cells with the minCMV>dsmCherry control construct.

### Immunofluorescent staining

#### Coimmunostaining metastases for CSCs, tumor cells and Ki67

5µm frozen sections were air-dried for 5 mins before fixing for 10 mins at room temperature in 4% paraformaldehyde. Post-fixation samples were washed, permeabilized with 0.5% Triton X-100 for 3 mins, washed again and blocked in PBS/10% BSA for 1h at room temperature. Blocked sections were incubated overnight at 4oC with the following primary antibodies diluted in PBS/10% BSA: Ki67 (proliferation), Abcam ab16667, 1:1000; mCherry (CSCs), Thermofisher M11217, 1:500; GFP (all tumor cells), Abcam ab6673, 1:500). After washing with PBS, sections were incubated with secondary antibodies conjugated to Alexa Fluor dyes (Invitrogen, Carlsbad, CA: Anti-Goat-IgG Alexa 488, Cat#A11055; Anti-rabbit-IgG Alexa 647 at 1:500 in PBS/10% BSA for 1 hour at room temperature, protected from light. After washing, tissues were counterstained with DAPI (300nM) and mounted with ProLong Gold Antifade Mountant (Thermo Fisher Scientific, Waltham, MA). Fluorescent images were acquired using a Zeiss LSM 880 confocal microscope (Carl Zeiss Microscopy GmbH, Jena, Germany) and 40x/1.3 oil DIC UV-IR M27 lens with a numerical aperture (NA) of 1.3. Image acquisition settings were standardized across all samples.

#### OPAL immunostaining metastases for tumor cells and YAP

Immunostaining for tumor cells and YAP in metastases was performed using formalin-fixed, paraffin-embedded (FFPE) metastatic mouse lung sections. The staining was performed on a Leica Biosystems Bond RX autostainer using the Bond Polymer Refine Kit (DS9800, Leica Biosystems), omitting the PostPrimary reagent, DAB, and Hematoxylin. Antigen retrieval was conducted using Citrate (Bond Epitope Retrieval 1). Sections were incubated with anti-YAP antibody (14074, Cell Signaling Technology; 1:200) for 30 min, followed by the Polymer reagent and OPAL Fluorophore 5705. The YAP antibody complex was stripped using Bond Epitope Retrieval 24. Sections were then incubated with anti-GFP antibody (ab183734, Abcam; 1:1200) for 30 min, followed by the Polymer reagent and OPAL Fluorophore 5206. Finally, sections were stained with DAPI and coverslipped using Prolong Gold AntiFade Reagent (Invitrogen). Images were captured using an Aperio Scanscope FL whole slide scanner. Quantitative image analyses were done using Halo software.

#### Immunofluorescent staining of cultured cells

LM2 cells were seeded into Ibidi 8-well chamber glass-bottom slides (Ibidi, Grafelfing, Germany) and cultured under standard conditions for 48h. Cells were then fixed in 4% PFA for 10 mins at room temperature, permeabilized with 0.5% TritonX-100 in PBS for 5 mins at room temperature and non-specific binding was blocked by incubating with 10% BSA in PBS for 1h at RT. Subsequent steps were as for the metastases above, but using the following primary antibodies (anti-YAP antibody (14074, Cell Signaling Technology; 1:200; anti-GFP antibody (ab183734, Abcam; 1:1000; GFP (all tumor cells), Abcam ab6673, 1:50) and secondary antibodies (Anti-Goat-IgG Alexa 488, Cat#A11055; Anti-Rat-IgG Alexa 568; Anti-rabbit-IgG Alexa 647).

### Image Analysis

#### HALO HighPlex

Quantitative image analysis for Ki67+ cells in large metastatic lesions was conducted using the HALO™ platform (Indica Labs,) with the HighPlex FL version 4.2.14 module. Metastatic lesions were manually delineated as regions of interest (ROIs) based on GFP fluorescence, which specifically marks cancer cells. To enable detailed marker quantification, cells within each ROI were segmented into nuclear and cytoplasmic subpopulations.

#### Image J

For assessment of CSCs and non CSCs in early lung colonization, captured images were processed to quantify the total tumor cells and CSCs in each metastatic lesion, using ImageJ to measure fluorescence intensity and count cells. GFP fluorescence or immunostaining identified all tumor cells. mCherry fluorescence, or immunostaining for mCherry or FLAG identified CSCs, and true positive signal was determined using parallel tissue samples with the minCMV>dsmCherry reporter to set the threshold.

#### YAP quantitation

For YAP-stained tissues, YAP was only quantified in tumor cells. Cells were segmented and quantified for YAP expression using a custom script written in Python (v. 3.10). Nuclear and cellular segmentation used the Cellpose deep-learning algorithm ^49^. DAPI intensity was used to segment nuclei using a custom model that was trained with the NIH HPC Biowulf cluster (http://hpc.nih.gov). A combined image of DAPI and GFP labeling was used for cellular segmentation using the pre-trained *cyto3* model. Cytoplasmic regions were created by subtracting nuclear ROIs from cellular ROIs. YAP intensity in nuclear and cytoplasmic regions were quantified, along with cellular SORE6>dsmCherry intensity.

### Statistical Analyses

Statistical analyses were performed in GraphPad Prism (version 10.1.1; GraphPad Software LLC, La Jolla, CA), with the exception of contingency table analyses which were performed in VassarStats (http://vassarstats.net). In general, non-parametric analyses were used for *in vivo*-generated data, and multiple comparison corrections were applied throughout where appropriate. The specific tests used are indicated in each figure. Statistical significance indicated by: *, p<0.05; **, p<0.01; ***, p<0.001; ****, p<0.0001.

## Data availability

All sequence data generated for this study can be accessed and downloaded from the Gene Expression Omnibus database using the accession numbers. The SORE6>dsmCherry and minCMV>dsmCherry prolentiviral plasmids have been deposited in the Addgene repository (addgene.org) with ID numbers 225665 and 225667.

