## Supplemental Information for "Hyper-responsiveness of cancer stem cells to microenvironmental cues controls metastasis and therapy response through YAP/TAZ/TEAD"

by

Tang *et al*.

### EXTENDED DATA

- Extended data figure legends
- Extended data figures
- Extended data table 1

#### EXTENDED DATA FIGURE LEGENDS

**Extended Data for Fig. 1| Breast cancer stem cells are enriched in the circulation and on arrival in the lung. a.** Schematic of fluorescent reporter and selectable marker cassette used in this study. The SORE6 enhancer element coupled to a minimal cytomegalovirus (CMV) promoter region responds to the presence of stem cell master transcription factors Sox2 and Oct4 or their paralogs. The mCherry is destabilized with a C-terminal degron from the ornithine decarboxylase (ODC) gene and has an N-terminal $FLAG_3$ tag for improved detection by immunohistochemistry or immunofluorescence. The selectable marker cassette has a C-terminally truncated version of the human CD19 protein driven by the constitutive SV40 promoter for FACS-based selection of transduced cells. A matched control lentivirus lacks the SORE6 enhancer but is otherwise identical. **b.** The half-life of the dsmCherry in LM2 cells *in vitro* was determined by following the decay in dsmCherry signal after the addition of cycloheximide at t=0 to block further protein synthesis. The signal from eGFP in the same cells is shown for comparison. **c.** Representative FACS profiles of transduced LM2 cells. **d.** Results of *in vivo* limiting dilution assays for tumor-initiating cells in sorted SORE6+ and SORE6- cell populations from 3 different TNBC cell lines. **e.** Summary of ELDA results for 6 different breast cancer models showing 8-20x enrichment of tumor initiating ability in SORE6+ vs SORE6- cells across all models.

**Extended Data for Fig. 2| CSCs and nonCSCs have d**ifferent population dynamics **in early lung colonization. a.** Number of surviving LM2 tumor cells in each subpopulation (CSC vs nonCSC) in the lung, normalized to time t=6h. The right y-axis indicates CSCs as % all tumor cells at each timepoint, where the injected tumor cell population had ~20% CSCs. Results are mean +/- SEM for n=3 mice/timepoint. **b.** Metastatic lesion size (# tumor cells/lesion) as a function of time after tail-vein injection of LM2 cells. Boxes show median +/- IQ range for pooled lesions from 1-3 mice/timepoint, for 2 separate experiments with different lung harvest times as indicated. Dunn's multiple comparisons test vs day 3. **c.** CSCs as a % total tumor cells/lung decreasing with time after tail-vein injection of LM2 cells. Results are mean +/- SD For n=3 mice/timepoint; Dunnett's multiple comparisons test v day 3 **d.** % CSCs in metastatic lesions of specified sizes as a function of time after tail-vein injection of LM2 cells. Results are mean +/- SEM for n=7-47 lesions/size bin/timepoint. **e.** # CSCs and nonCSCs per lesion as a function of metastatic lesion size in lungs of mice bearing orthotopic LM2 tumors at t=8 weeks after tumor cell implantation. **f.** % CSCs in lesion as a function of increasing metastatic lesion size in lungs from mice bearing 4T1 orthotopic primary tumors at t= 28 and 35 days after tumor cell implantation. Red lines on violin plots indicate medians; dotted lines are quartiles for 7-43 lesions/size bin, pooled from 4 mice. ANOVA **g,h.** Number of CSCs and nonCSCs/lesion as a function of lesion size for very small lesions in LM2 orthotopic model (**g**), or 4T1 orthotopic model (**h**). Results are mean +/- SD for 1-29 lesions/size bin. **i.** Number of metastases/lung at day 21 in mice injected with sorted SORE6+ or SORE6- 4T1 cells via tailvein on day 0. Results are median +/- IQ range. Mann-Whitney test. **j.** Metastatic lesions without CSCs as a function of lesion size in mice injected with unsorted LM2 cells via tail vein. Results are pooled from 5 mice. The number of lesions evaluated in each size bin is given on the graph. All lesions >= 32 cells in size had detectable CSCs.

**Extended Data for Fig. 3| Early inhibition of CSC population expansion is density-dependent and driven by increased CSC differentiation. a.** Images of LM2 culture composition at time of peak CSC plateau for cultures with different input % CSCs as indicated. **b**. % CSCs proliferating at different times in LM2 and MCF10Ca1h cultures as assessed by single cell fate mapping. Early = 36-60h and Late = 60-84h after cell plating. Data replotted from **Fig. 3n**. Fisher Exact test for division-dependent vs other events.

**Extended Data for Fig. 4| CSCs are hyper-responsive to microenvironmental inputs. a,b.** Incucyte Live Cell imaging to assess the effect of the FAK inhibitor defactinib on CSCs and nonCSCs in unsorted 4T1 **(a)** and SUM159 **(b)** cultures. Arrow indicates time of addition of drug (24h). Results are mean +/- SD, n=3. **c,d.** Incucyte Live Cell imaging to assess effect of medium refreshment on CSCs and nonCSCs in unsorted 4T1 **(c)** and SUM159 **(d)** cultures. Arrow indicates time of medium change.

**Extended Data for Fig. 5| Multiple microenvironmental inputs to the CSC sensor converge on YAP/TAZ a.** Representative GSEA plots for enrichment of stem-related genesets in LM2 CSCs *in vitro*. **b.** Immunofluorescent staining to assess YAP localization in low and high density LM2 cultures. Results are from n=197 cells (low density culture) or n=1220 cells (high density culture). Solid line shows median and dotted lines interquartile ranges; Mann-Whitney test. **c.** Representative immunofluorescent image of YAP localization in tumor cells in small and large LM2 lung metastases. Nuclear localization indicates active YAP. Line shows median for 6-7 metastatic lesions per size bin. Mann-Whitney test. **d.** Western blot of YAP phosphorylation status in response to indicated culture conditions. CM indicates treatment with conditioned medium from high density culture. Phospho-YAP is the inactive form. **e.** Quantitation of YAP localization by immunofluorescence in LM2 cultures following treatment with the ROCK inhibitor Y27632. n=330 (Con) and n=400 (Y7632); Mann-Whitney test. **f.** RNA expression in unsorted LM2 cultures *in vitro* of genes relating to LPA biosynthesis and response. **g,h** Incucyte live cell imaging for effect of LPAR inhibitor BMS5986020 (BMS) **(g)** or LPA **(h)** on CSCs and nonCSCs in unsorted 4T1 cultures. Arrows indicate time of treatment. Mean +/- SD (n=3).

**Extended Data for Fig. 7| Targeting CSC sensor nodes reduces chemotherapy-driven enrichment of CSCs in metastasis. a.** Schematic for treatment with GNE7823 and paclitaxel in the LM2 model. **b.** Effect of TEAD inhibitor GNE7832 and paclitaxel on metastasis number in the LM2 model. Results are median +/- IQ range, for n=7 mice/group; Dunn's multiple comparison test vs PAC. **c.** Schematic for treatment of LM2 model with LPAR inhibitor BMS5986020 (BMS) and Paclitaxel (Pac). **d.** Median lesion size (# tumor cells) in lungs from mice treated with LPAR inhibitor (BMS) and paclitaxel (Pac). Results are median +/- IQ range, n=4-6 mice/group; Dunn's multiple comparisons test vs CON. **e.** Treatment effect on number of CSCs/mouse lung. Median +/- IQ range, n=4-6 mice/group; Dunn's multiple comparisons test vs CON. **f.** Treatment effect on %CSCs/lesion as a function of metastatic lesion size. Data pooled across all mice/treatment group, n=4-6 mice/group; 3-58 lesions/size bin/treatment. Dunn's multiple comparisons test. Only significant differences shown. **g.** Effect of treatment on % metastatic lesions with no CSCs. Data pooled across all mice/treatment group, n=4-6 mice/group; total lesion numbers per treatment group indicated on graph. Fisher exact test for CON v BMS and PAC v PAC+ BMS.

#### Extended Data Fig. 1 | SORE6 reporter design and validation

**a**

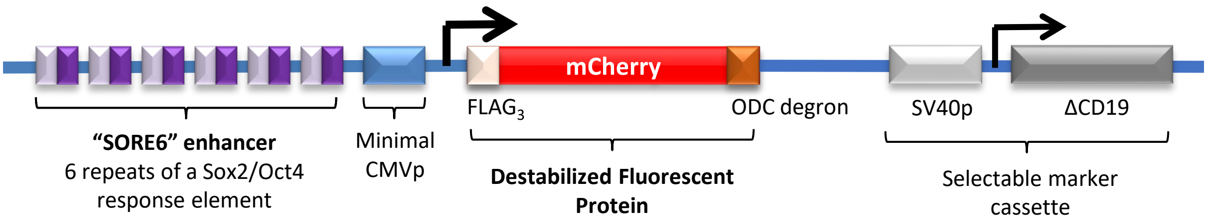

**b**

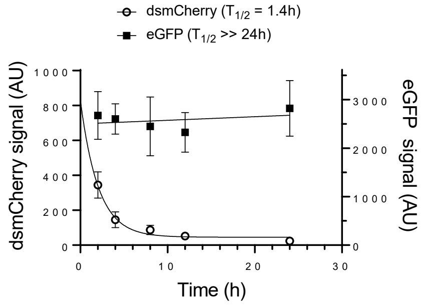

**c**

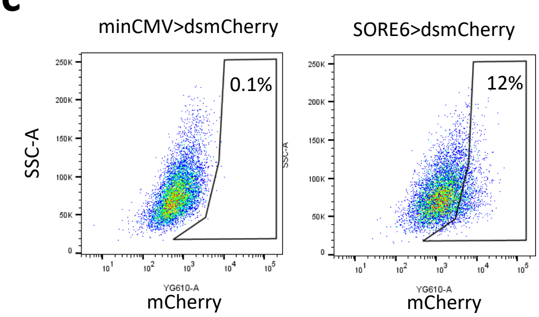

**d**

| Cells injected | | Tumor assessment time (days) | Cell innoculum and number of tumors formed 10000 | 5000 | 2000 | 1000 | 500 | 250 | 200 | 100 | CSC frequency |
|---|---|---|---|---|---|---|---|---|---|---|---|
| MDAMB231-LM2 | SORE6+ | 62 | - | - | 1/6 | - | 4/6 | - | - | 2/4 | 1/1868 |
| MDAMB231-LM2 | SORE6- | 62 | - | - | 1/6 | - | 0/6 | - | - | 0/6 | 1/14577 |
| SUM149 | SORE6+ | 75 | 4/6 | 5/6 | - | 3/6 | - | - | - | 0/6 | 1/4442 |
| SUM149 | SORE6- | 75 | 0/6 | 2/6 | - | 0/6 | - | - | - | 0/6 | 1/45754 |
| SUM159 | SORE6+ | 70 | - | 6/6 | - | 6/6 | 4/6 | 4/6 | - | - | 1/305 |
| SUM159 | SORE6- | 70 | - | 1/6 | - | 3/6 | 3/6 | 2/4 | - | - | 1/3821 |
| 4T1 | SORE6+ | 21 | 10/10 | - | - | 10/10 | - | - | 7/10 | - | 1/163 |
| 4T1 | SORE6- | 21 | 9/10 | - | - | 6/10 | - | - | 0/10 | - | 1/2737 |

**e**

| MODEL | Breast cancer subtype | Species | Enrichment factor | Chi-sq P-value | Source data |
|---|---|---|---|---|---|
| MCF7 | ER+ | Human | **>21** | 1.2e-09 | Tang et al. |
| MCF10Ca1h | Ras-transformed HMEC | Human | **20** | 6.9e-08 | Tang et al. |
| SUM159 | TNBC | Human | **12.5** | 4.4e-08 | This manuscript |
| SUM149 | TNBC | Human | **10.3** | 2.2e-04 | This manuscript |
| MDAMB231 (parental ATCC) | TNBC | Human | **9** | 4.5e-08 | Sharma et al. |
| MDAMB231-LM2 | TNBC | Human | **7.8** | 0.017 | This manuscript |
| 4T1 | TNBC | Mouse | **17** | 8.7e-09 | This manuscript |

#### Extended Data Fig. 2 | CSCs and nonCSCs have different population dynamics in early lung colonization

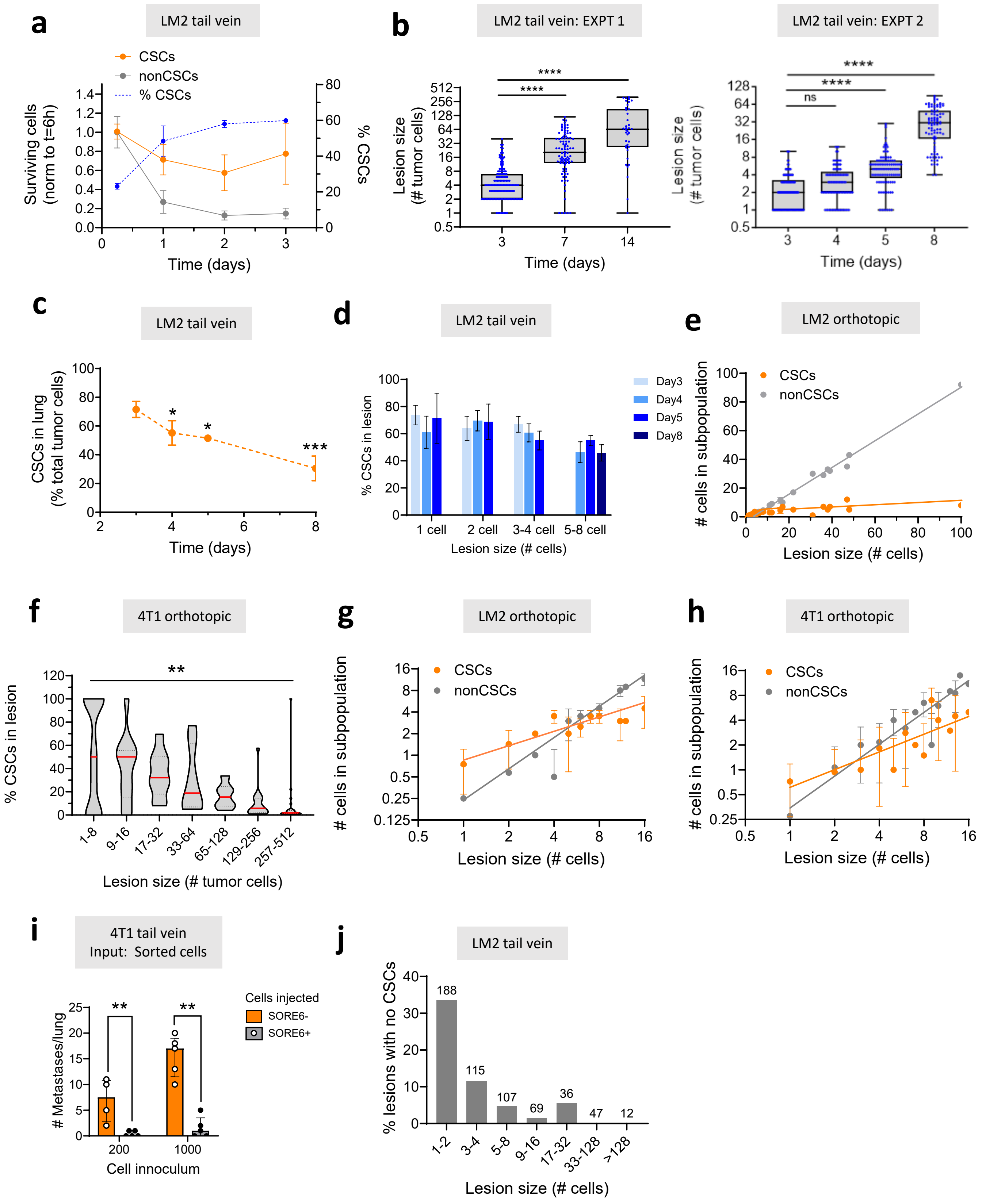

**Extended Data for Fig. 3| Early inhibition of CSC population expansion is density-dependent and driven by increased CSC differentiation**

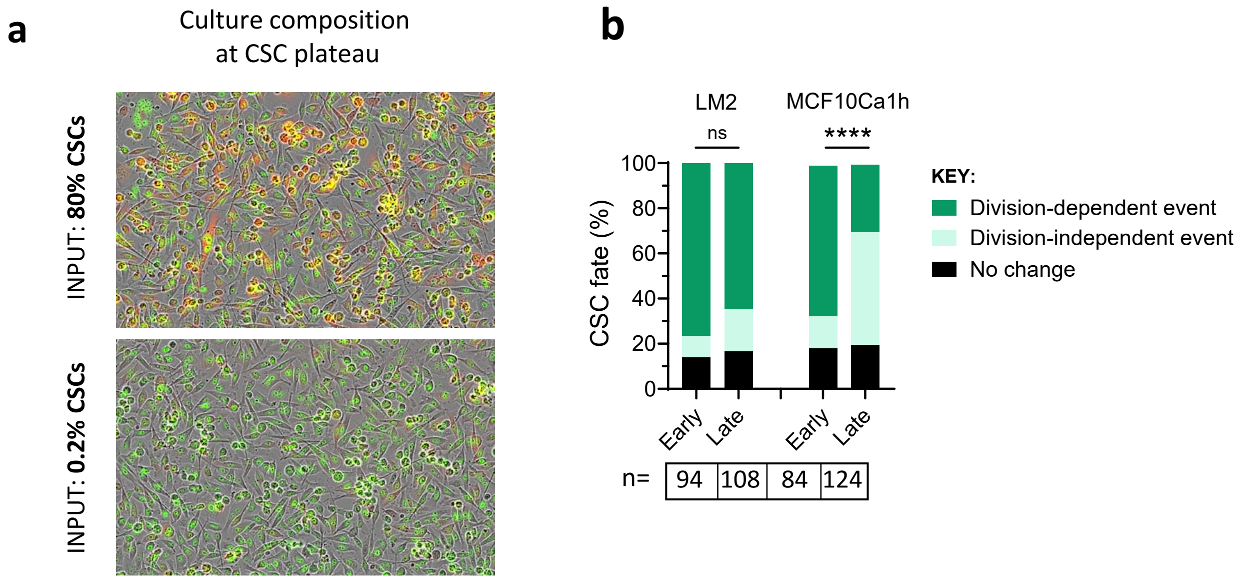

#### Extended Data for Fig. 4| CSCs are hyper-responsive to microenvironmental inputs

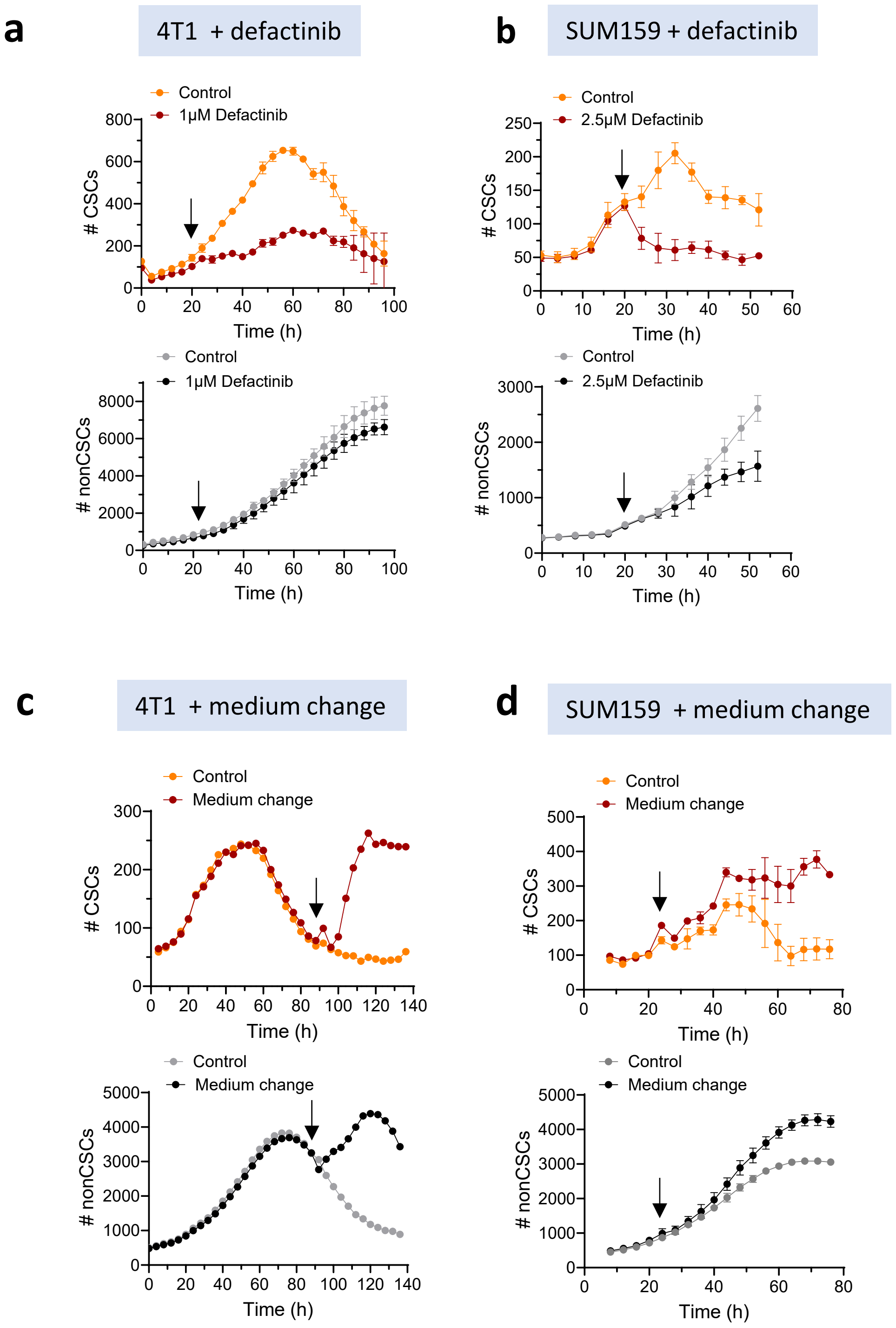

#### Extended Data for Fig. 5| Multiple microenvironmental inputs to the CSC sensor converge on YAP/TAZ

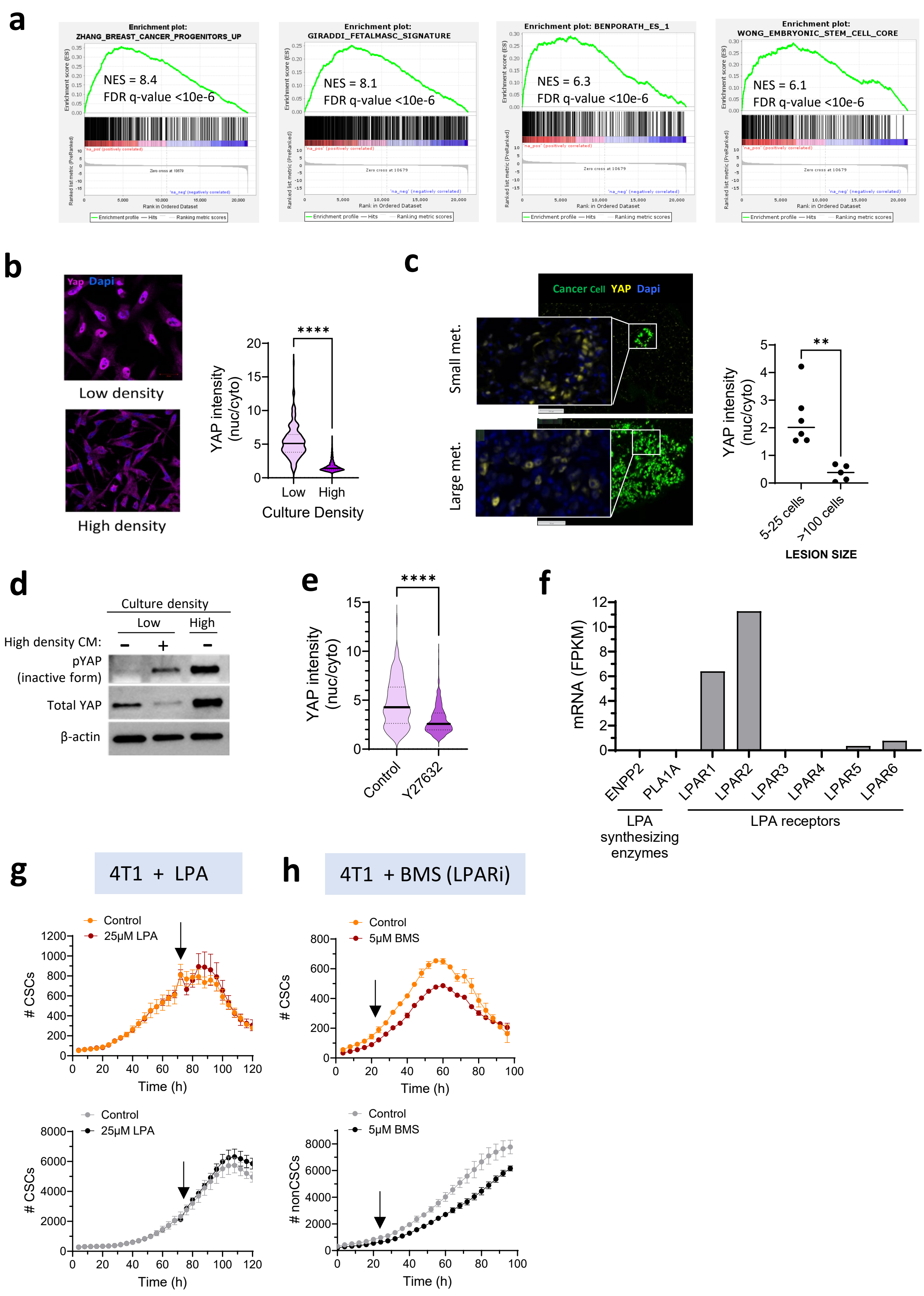

**Extended Data for Fig. 7| Targeting CSC sensor nodes reduces chemotherapy-driven enrichment of CSCs in metastasis**

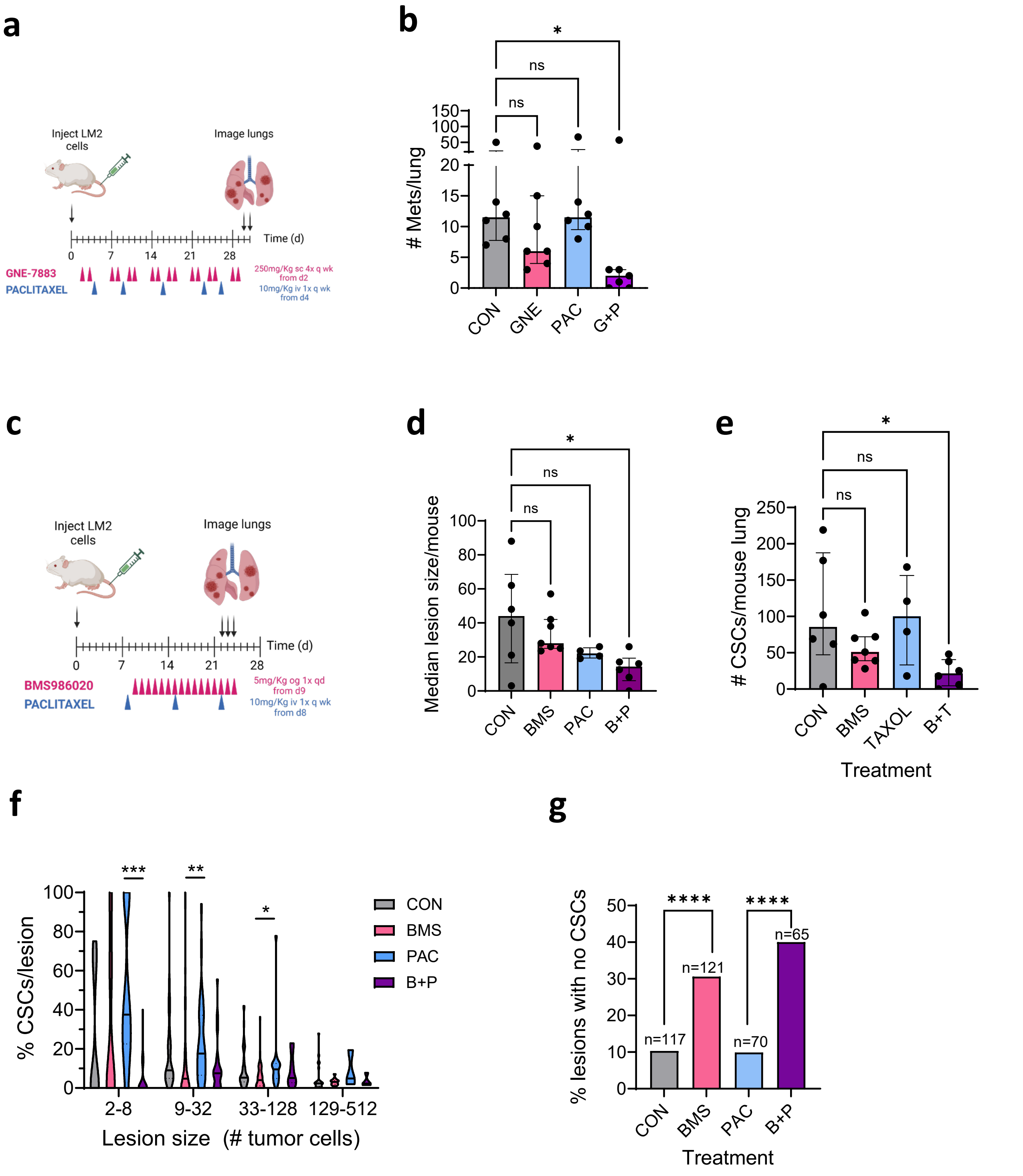

**Extended Data Table 1| Effect of pharmacologic modulation of upstream inputs to YAP/TAZ node on CSC dynamics**

| Biological inputs | Molecular target | Pharmacologic agent | Predicted effect of drug on YAP/TAZ | Observed Effect on CSCs | | |
|---|---|---|---|---|---|---|
| | | | | LM2 | SUM159 | 4T1 |
| Cell-cell contact/ polarity | LATS kinase | TRULI | Stimulation | ++ | +++ | ++ |
| Integrin-mediated cell attachment and spreading | FAK | Defactinib | Inhibition | - - | - - | - - - |
| | F-actin | Cytochalasin D | Inhibition | - - | - | - - |
| Bioactive lipids | LPAR | BMS986020/ AM966 | Inhibition | - - - | - - | - |
| Bioactive lipids and/or ECM attachment | ROCK | Y27632/ H1152/ Fasudil | Inhibition | - - - | - - - | - - - |
| Cellular energy stress | mTORC | Torkinib | Inhibition | - | - | - |
| | AMPK | AICAR*/ MK3903* | Inhibition | - | - - | - |

*Pharmacologic activator

**Key:**

**Stimulation:** + (little or no effect); ++ (moderate effect); +++ (strong effect)

**Inhibition:** - (little or no effect); - - (moderate effect); - - - (strong effect)

**Extended Data Table 1| Effect of pharmacologic modulation of upstream inputs to YAP/TAZ node on CSC dynamics** Pharmacologic agents targeting potential upstream inputs to the YAP/TAZ node were tested at multiple concentrations across three breast cancer models for selective effects on the CSC population. CSC and nonCSCs were assessed by Incucyte live cell imaging in unsorted cell cultures. Scores reflect the relative effect size for the agent specifically on the CSCs. Some agents (e.g. Torkinib) affected both CSCs and nonCSCs, but had a greater effect on CSCs. ROCK lies downstream of both bioactive lipid signaling and integrin activation by ECM attachment.
